# Transcriptomic and Proteomic Analyses of a New Cytoplasmic Male Sterile Line with wild *Gossypium bickii.* Genetic Background

**DOI:** 10.1101/2020.02.11.943464

**Authors:** Haiyan Zhao, Jianshe Wang, Yunfang Qu, Renhai Peng, Richard Odongo Magwanga, Fang Liu, Jinling Huang

## Abstract

Cotton is an important fiber crop but has serious effects of heterosis, in which cytoplasmic male sterility (CMS) being the major cause of heterosis in plants. However, there are no studies done on CMS Yamian A in cotton with the genetic background of the Australian wild *Gossypium bickii.* Transcriptomic and proteomic results showed that UDP-glucosyltransferase - in the nucleus, 60S ribosomal protein L13a- in the cytoplasm, ribulose-1,5-bisphosphate carboxylase/oxygenase - in the chloroplast, glutathione S-transferase - in the cytoplasm, and ATP synthase F1 subunit 1 - in the mitochondrion were upregulated; while low molecular weight heat shock protein - in the chloroplast and ATP synthase D chain- in the mitochondrion were down-regulated expression at the microspore abortion stage of Yamian A. We constructed an interaction network and this study provides a comprehensive understanding of the mechanism of CMS in cotton by use of in Yamian A, with wild cotton genetic background.

## INTRODUCTION

Cotton is an important cash crop with high-quality fiber, edible oil, and protein which is mainly used as animal feeds^(1)^. Heterosis in cotton is quite apparent and has been widely used in yield quality, and resistance studies in cotton^(2)^. The adoption of the production of hybrid seeds is the most important link among some links of cotton heterosis use. At present, the castration in the production of hybrid seeds often relies on hand emasculation and male-sterile lines including chemically induced male sterility (IMS), genic male sterility (GMS), and cytoplasmic male sterility (CMS)^(3)^. The production practice showed that CMS is an effective way of heterosis utilization in crop and widely used to produce hybrid seeds, because it eliminates the need for artificial emasculation, saves manpower and material resources, enhance the purity of hybrid seeds and increases the output of crops^(3, 4)^. All over the world, cotton studies started in the 1960s, since then a number of germplasms have been developed, such as *G. arboreum* L, *G. harknessii* Brandegee, *G. trilobum* (DC.) Skov., *G. hirsutum*, *G. barbadense* L., among others. However, there is no report on CMS in cotton with genetic background of Australian wild *Gossypium bickii*, which has been reported despite the enormous effects of heterosis in cotton germplasm development.

In recent years, advancement in molecular technology, has enabled breeders and molecular researchers to identify various plant transcription factors, genes and explore protein expression at the translational, transcriptome and proteome level, such as the CMS studies of the Chinese cabbage^(5)^, turnip^(6)^, *Cucumis melo* L.^(7)^, cotton ^(8, 9)^, rice^(10)^, *Brassica napus* L.^(11)^. Transcriptomic analysis in cotton (CMS-D8) revealed that the reactive oxygen species (ROS) were released from mitochondria and served as important signal molecules in the nucleus, causing the formation of abnormal tapetum^(8)^. Proteome analyses in cotton indicated that the differentially expressed proteins (DEPs) mainly involved in pyruvate, carbohydrate and fatty acid metabolism had been identified between the male-sterile line 1355A and the male-fertile line 1355B^(9)^. Integrated analysis of the transcriptome and proteome can afford a complete picture in regard to the molecular mechanism of CMS, and have been reported in Chinese cabbage^(12)^, *Brassica napus*^(13)^, Pepper^(14)^, *Citrus suavissima*^(15)^ involved CMS studies. The conjoint analysis of the transcriptome and proteome in Shaan2A CMS and its maintainer line revealed that the sterility gene from mitochondrion might suppress the expression of relevant transcription factor genes in the nucleus, affecting early anther development^(13)^. There have been relatively few studies of the conjoint analysis of transcriptomic and proteomic changes in cotton CMS at the moment.

Yamian A was identified by the cotton breeding group of Shanxi Agricultural University, as a new and stable cytoplasmic male sterile line derived from the triple hybrids of *Gossypium bickii*, *Gossypium arboreum* and *Gossypium hirsutum* Linn^(16)^. The male sterility mechanism of Yamian A CMS is unclear. In the current study, the conjoint analysis of the transcriptomic, proteomic and early cytological, physiological and biochemical, were first performed between the Yamian A and its maintainer Yamian B, to reveal the mechanism of Yamian A CMS. We aimed to identify differentially expressed genes and proteins at different development stages of anthers, and discuss the relationship between these differentially expressed genes and proteins and male sterility in Yamian A CMS, and explore the possible effects on microspore abortion of Yamian A CMS. These study results will offer new insights into the molecular mechanism of Yamian A CMS and improve our understanding of male sterility in cotton.

## MATERIALS and METHODS

### Plant materials

The cotton CMS line Yamian A (YA-CMS) and its maintainer Yamian B (YB) were planted in the experimental field of Shanxi Agricultural University, Taigu, Shanxi, China, during the natural growing season. According to Cytological observation, the cotton flower buds of the CMS line Yamian A and its maintainer Yamian B was divided into seven consecutive grades (Table S1). Stage 1: the buds were the normal development and before microspore abortion stage; Stage2, 3 and 4: the buds were the fertility transformation and middle microspore abortion stage, stage 2, 3 were the key stage of pollen abortion; Stage5, 6 and 7: the buds were entirely abortive and after microspore abortion stage^(16)^. At the anthesis, the buds of before, middle and after microspore abortion stage of Yamian A and Yamian B were collected for the transcriptome research; the buds of the key stage of pollen abortion (Stage2, 3) were collected for the proteomics research; the buds of seven different development periods respectively were used to do expression analysis by quantitative real-time PCR (qRT-PCR). An individual hybrid dynamic sampling method was used in the sampling process to ensure that each sample has the same genetic background and the growth period.

### Transcriptome analysis

The total RNA of the buds collected for the transcriptome research was extracted by the EASYspin Plus Plant RNA Kit RN37 (Aidlab Biotechnology) and cDNA synthesis by the M-MLV RTase cDNA Synthesis Kit (TaKaRa Company). cDNA amplified fragment length polymorphism (cDNA-AFLP) analysis was performed and made a little change as described previously^(17)^. The differentially expressed band’s sequences were analyzed with the DNASTAR software, BLAST instrument at the National Center for Biotechnology Information (http://www.ncbi.nlm.nih.gov/blast) program, GO (Gene Ontology) (http://www.blast2go.org/) and KEGG (http://www.genome.jp/kegg/).

### Proteomics Analysis

Protein isolate, 2-DE, image analysis, tryptic digestion and identification of differentially expressed proteins were performed as described previously with some modifications^(18)^. The Uniprot Knowledgebase (http://www.uniprot.org/) with the NCBI (http://www.ncbi.nlm.nih.gov/), and the Gene Ontology (GO) database were used to classify identified proteins into the specific biological process and cellular component. The STRING 9.05 (http://string-db.org/) was used to do enrichment analysis of molecular function and metabolic pathways with Arabidopsis thaliana as an analysis reference database for GO and the Kyoto encyclopedia of genes and genomes (KEGG). The STRING 9.05 (http://string-db.org/) was also used to do a protein-protein interaction network of differential proteins with Arabidopsis thaliana as reference species.

### qRT-PCR

Total RNAs extraction, reverse transcription and qRT-PCR from the buds of seven different development periods respectively of both the fertile and sterile plants were performed respectively using EASYspin Plus Plant RNA Kit RN09 (Aidlab Biotechnology), PrimeScript® RT Master Mix Perfect Real-Time and DRR820ASYBR® Premix Ex Taq™ II (Tli RNaseH Plus) (TaKaRa), according to the manufacturer’s instructions. Each sample was performed in three biological replicates. The relative expression of the target genes was calculated with the 2^-△△Ct^ method^(19)^. Primers for qRT-PCR analysis are shown in Supplementary Table S2. There were three biological with three technical replicates per sample.

## RESULTS

### Transcriptome analysis Expression type of the differently expressed fragments

The cDNA-AFLP analysis was used to do transcriptome research between Yamian A and Yamian B with the buds of before, middle, and after microspore abortion stage. A total of 256 primer combinations were screened, 134 of them produced 550 differently expressed fragments. These differently expressed fragments were not only the difference of quantity but also the distinction of the quality (Figures S1). Expression type of the differently expressed fragments in the buds between Yamian A and Yamian B mainly included fifteen independent sets (Table S3); fragments detected at only one of the three stages in Yamian A or Yamian B (Type 1-6), especially the band number of type 2 was the most among all the types; fragments detected at any two of the three stages in Yamian A or Yamian B (Type 7-11), for example, the 12 fragments of type 7 were detected at the buds of before and middle microspore abortion stage in Yamian A; fragments detected at one or any two of the three stages in Yamian A and Yamian B (Type 12-15), for example, the 20 fragments of type 12 were detected at the buds of middle microspore abortion stage in Yamian A and Yamian B.

### Homology analysis of differentially expressed fragments

132 Transcript-derived fragments (TDFs) selected from 550 differently expressed fragments were recycled, cloned, and sequenced, 99 fragments were produced readable sequences ultimately (Table 1). The sizes of 99 fragments were concentrated between 19 to 500 bp. Sequence alignment of these 99 TDFs revealed that 31 showed homology to genes with known functions, whereas one did not show homology to other sequences, and 67 displayed identity with unknown proteins. Sequence analysis indicated that some different TDFS come from different primer combinations were searched for the same homologous sequences, such as homologous sequences of T26 and T27 both were putative UDP-glucosyl transferase of *Ricinus communis*.

**Table 1.**
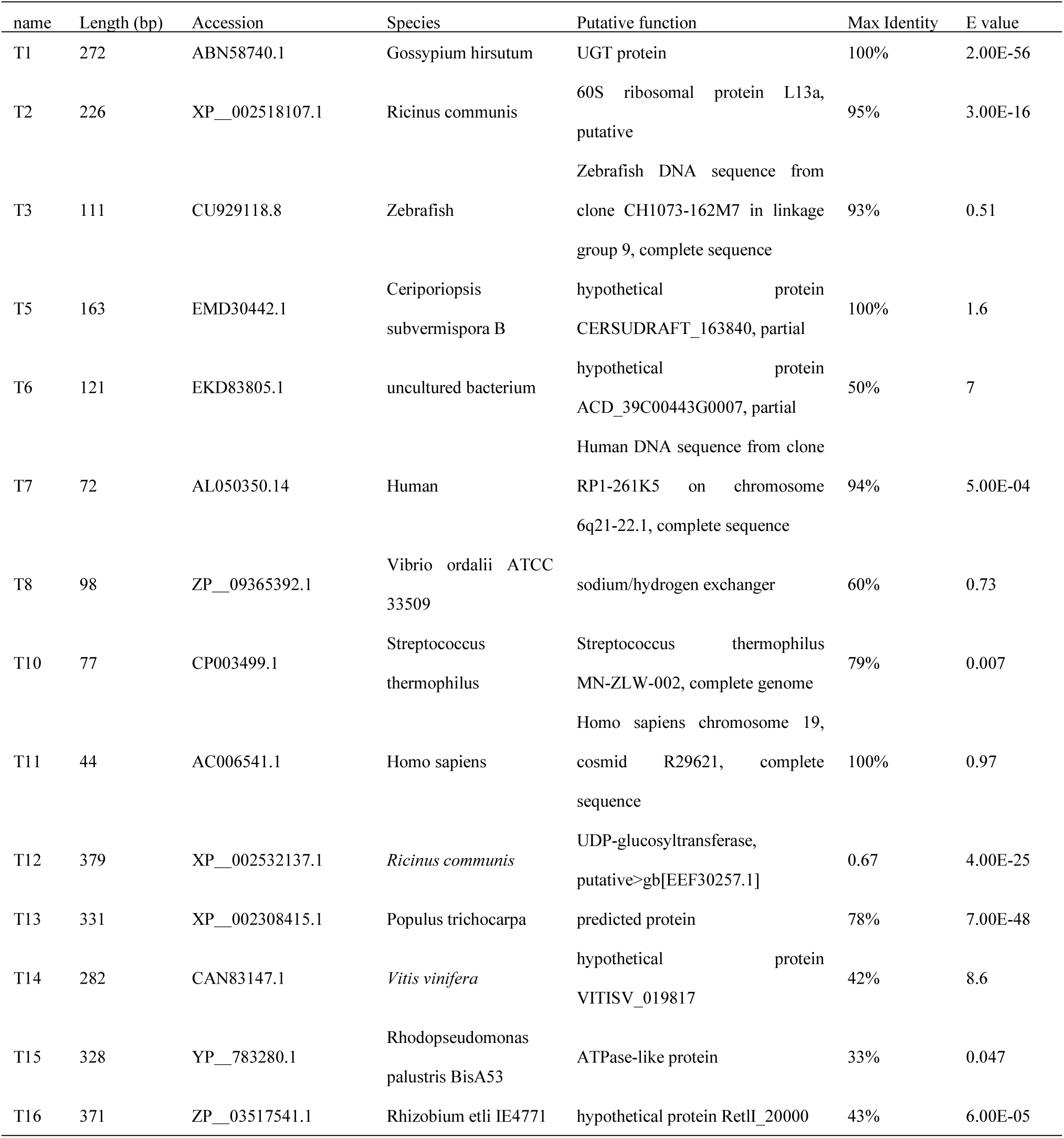
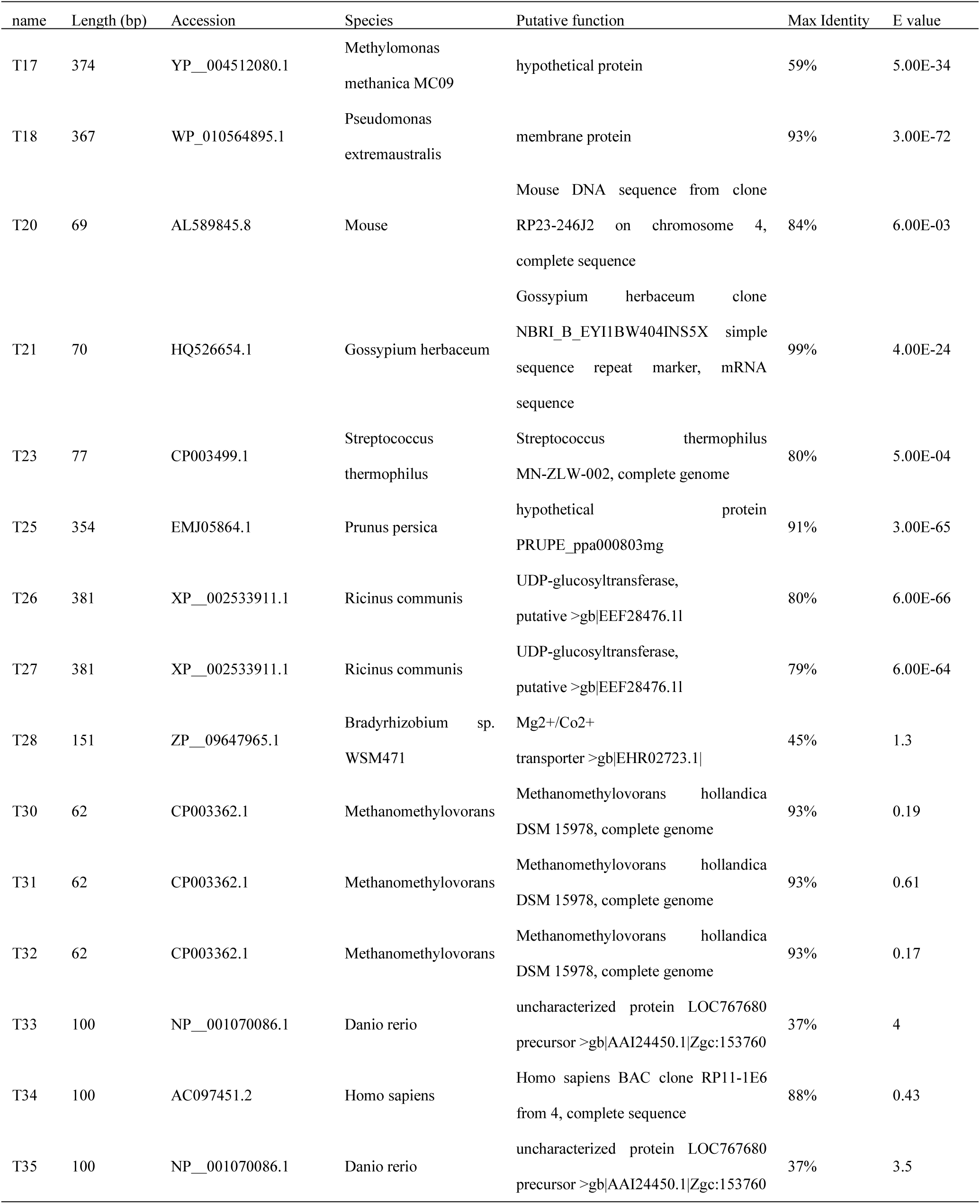
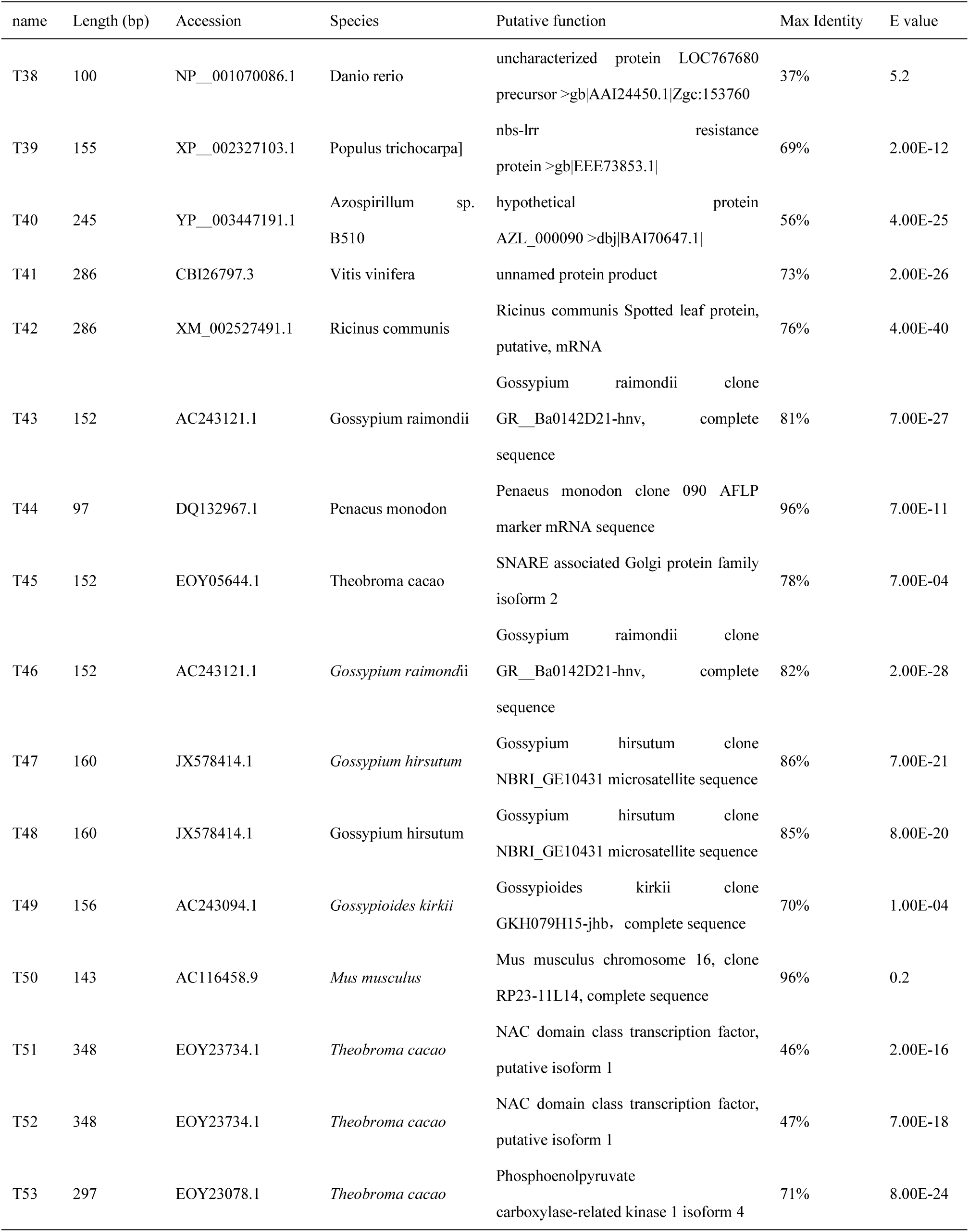
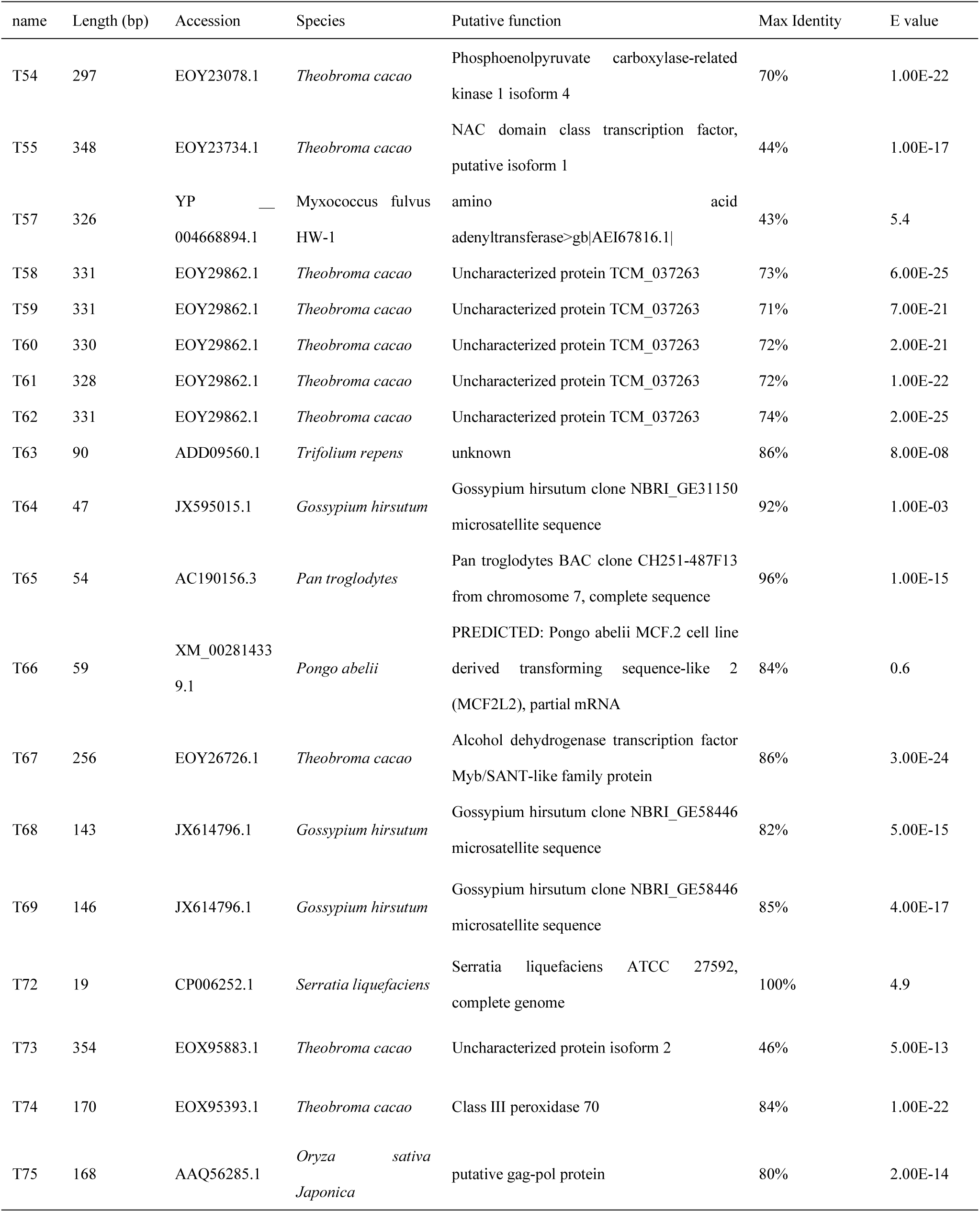
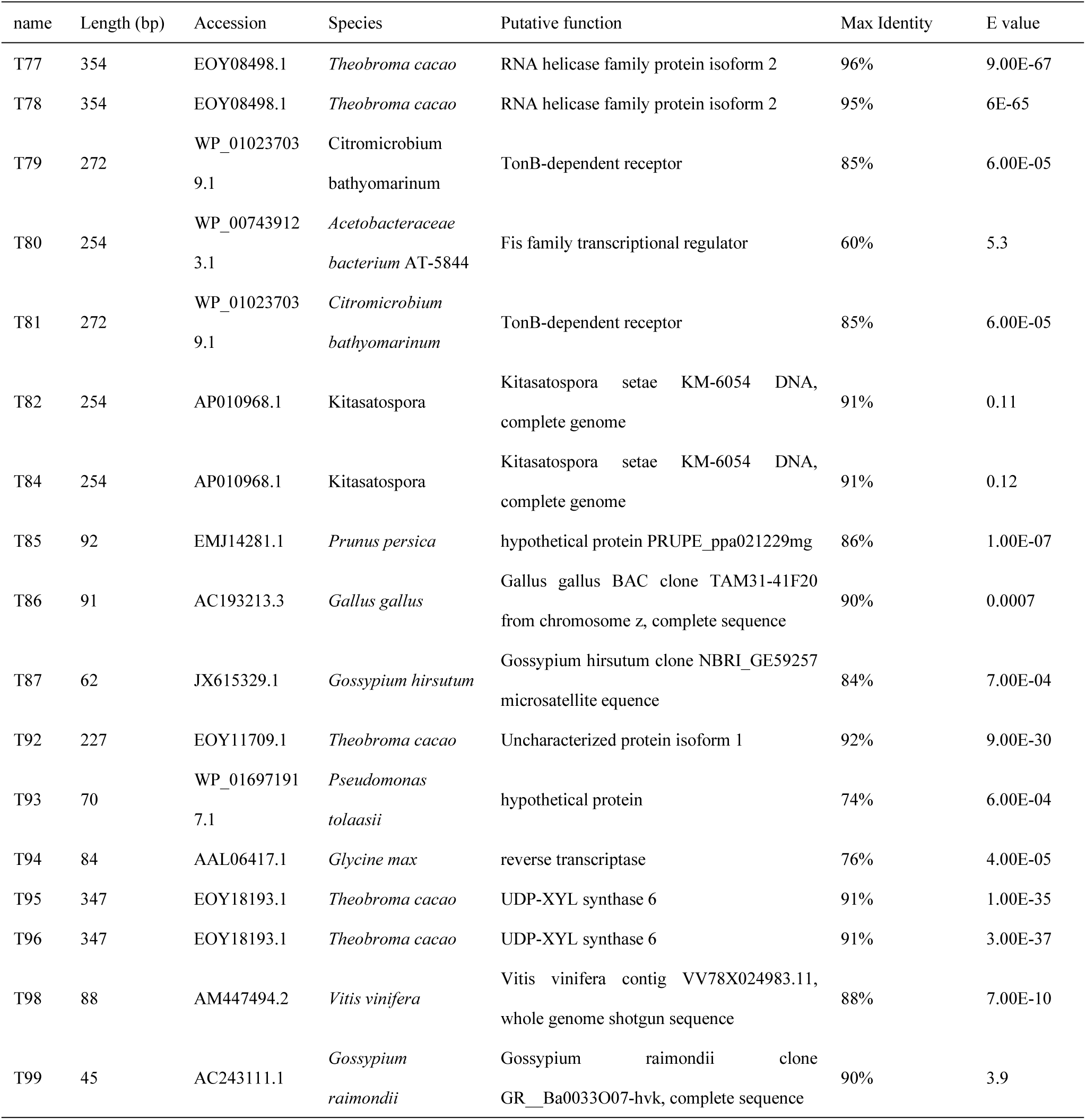
Homology analysis of TDFs sequences on cDNA - AFLP in GenBank.

| name | Length (bp) | Accession | Species | Putative function | Max Identity | E value |
| --- | --- | --- | --- | --- | --- | --- |
| T1 | 272 | ABN58740.1 | Gossypium hirsutum | UGT protein | 100% | 2.00E-56 |
| T2 | 226 | XP__002518107.1 | Ricinus communis | 60S ribosomal protein L13a,<br>putative | 95% | 3.00E-16 |
| T3 | 111 | CU929118.8 | Zebrafish | Zebrafish DNA sequence from<br>clone CH1073-162M7 in linkage<br>group 9, complete sequence | 93% | 0.51 |
| T5 | 163 | EMD30442.1 | Ceriporiopsis<br>subvermispora B | hypothetical protein<br>CERSUDRAFT_163840, partial | 100% | 1.6 |
| T6 | 121 | EKD83805.1 | uncultured bacterium | hypothetical protein<br>ACD_39C00443G0007, partial | 50% | 7 |
| T7 | 72 | AL050350.14 | Human | Human DNA sequence from clone<br>RP1-261K5 on chromosome<br>6q21-22.1, complete sequence | 94% | 5.00E-04 |
| T8 | 98 | ZP__09365392.1 | Vibrio ordalii ATCC<br>33509 | sodium/hydrogen exchanger | 60% | 0.73 |
| T10 | 77 | CP003499.1 | Streptococcus<br>thermophilus | Streptococcus thermophilus<br>MN-ZLW-002, complete genome | 79% | 0.007 |
| T11 | 44 | AC006541.1 | Homo sapiens | Homo sapiens chromosome 19,<br>cosmid R29621, complete<br>sequence | 100% | 0.97 |
| T12 | 379 | XP__002532137.1 | Ricinus communis | UDP-glucosyltransferase,<br>putative>gb[EEF30257.1] | 0.67 | 4.00E-25 |
| T13 | 331 | XP__002308415.1 | Populus trichocarpa | predicted protein | 78% | 7.00E-48 |
| T14 | 282 | CAN83147.1 | Vitis vinifera | hypothetical protein<br>VITISV_019817 | 42% | 8.6 |
| T15 | 328 | YP__783280.1 | Rhodopseudomonas<br>palustris BisA53 | ATPase-like protein | 33% | 0.047 |
| T16 | 371 | ZP__03517541.1 | Rhizobium etli IE4771 | hypothetical protein RetI_20000 | 43% | 6.00E-05 |
| T17 | 374 | YP__004512080.1 | Methylomonas<br>methanica MC09 | hypothetical protein | 59% | 5.00E-34 |
| T18 | 367 | WP_010564895.1 | Pseudomonas<br>extremaustralis | membrane protein | 93% | 3.00E-72 |
| T20 | 69 | AL589845.8 | Mouse | Mouse DNA sequence from clone<br>RP23-246J2 on chromosome 4,<br>complete sequence | 84% | 6.00E-03 |
| T21 | 70 | HQ526654.1 | Gossypium herbaceum | Gossypium herbaceum clone<br>NBRI_B_EYI1BW404INS5X simple<br>sequence repeat marker, mRNA<br>sequence | 99% | 4.00E-24 |
| T23 | 77 | CP003499.1 | Streptococcus<br>thermophilus | Streptococcus thermophilus<br>MN-ZLW-002, complete genome | 80% | 5.00E-04 |
| T25 | 354 | EMJ05864.1 | Prunus persica | hypothetical protein<br>PRUPE_ppa000803mg | 91% | 3.00E-65 |
| T26 | 381 | XP__002533911.1 | Ricinus communis | UDP-glucosyltransferase,<br>putative >gb\|EEF28476.1\| | 80% | 6.00E-66 |
| T27 | 381 | XP__002533911.1 | Ricinus communis | UDP-glucosyltransferase,<br>putative >gb\|EEF28476.1\| | 79% | 6.00E-64 |
| T28 | 151 | ZP__09647965.1 | Bradyrhizobium sp.<br>WSM471 | Mg<sup>2+</sup>/Co<sup>2+</sup><br>transporter >gb\|EHR02723.1\| | 45% | 1.3 |
| T30 | 62 | CP003362.1 | Methanomethylovorans | Methanomethylovorans hollandica<br>DSM 15978, complete genome | 93% | 0.19 |
| T31 | 62 | CP003362.1 | Methanomethylovorans | Methanomethylovorans hollandica<br>DSM 15978, complete genome | 93% | 0.61 |
| T32 | 62 | CP003362.1 | Methanomethylovorans | Methanomethylovorans hollandica<br>DSM 15978, complete genome | 93% | 0.17 |
| T33 | 100 | NP__001070086.1 | Danio rerio | uncharacterized protein LOC767680<br>precursor >gb\|AAI24450.1\|Zgc:153760 | 37% | 4 |
| T34 | 100 | AC097451.2 | Homo sapiens | Homo sapiens BAC clone RP11-1E6<br>from 4, complete sequence | 88% | 0.43 |
| T35 | 100 | NP__001070086.1 | Danio rerio | uncharacterized protein LOC767680<br>precursor >gb\|AAI24450.1\|Zgc:153760 | 37% | 3.5 |
| T38 | 100 | NP__001070086.1 | Danio rerio | uncharacterized protein LOC767680 precursor >gb\|AAI24450.1\|Zgc:153760 | 37% | 5.2 |
| T39 | 155 | XP__002327103.1 | Populus trichocarpa] | nbs-lrr resistance protein >gb\|EEE73853.1\| | 69% | 2.00E-12 |
| T40 | 245 | YP__003447191.1 | Azospirillum sp. B510 | hypothetical protein AZL_000090 >dbj\|BAI70647.1\| | 56% | 4.00E-25 |
| T41 | 286 | CBI26797.3 | Vitis vinifera | unnamed protein product | 73% | 2.00E-26 |
| T42 | 286 | XM_002527491.1 | Ricinus communis | Ricinus communis Spotted leaf protein, putative, mRNA | 76% | 4.00E-40 |
| T43 | 152 | AC243121.1 | Gossypium raimondii | Gossypium raimondii clone GR__Ba0142D21-hnv, complete sequence | 81% | 7.00E-27 |
| T44 | 97 | DQ132967.1 | Penaeus monodon | Penaeus monodon clone 090 AFLP marker mRNA sequence | 96% | 7.00E-11 |
| T45 | 152 | EOY05644.1 | Theobroma cacao | SNARE associated Golgi protein family isoform 2 | 78% | 7.00E-04 |
| T46 | 152 | AC243121.1 | *Gossypium raimondii* | Gossypium raimondii clone GR__Ba0142D21-hnv, complete sequence | 82% | 2.00E-28 |
| T47 | 160 | JX578414.1 | *Gossypium hirsutum* | Gossypium hirsutum clone NBRI_GE10431 microsatellite sequence | 86% | 7.00E-21 |
| T48 | 160 | JX578414.1 | *Gossypium hirsutum* | Gossypium hirsutum clone NBRI_GE10431 microsatellite sequence | 85% | 8.00E-20 |
| T49 | 156 | AC243094.1 | *Gossypoides kirkii* | Gossypoides kirkii clone GKH079H15-jhb, complete sequence | 70% | 1.00E-04 |
| T50 | 143 | AC116458.9 | *Mus musculus* | Mus musculus chromosome 16, clone RP23-11L14, complete sequence | 96% | 0.2 |
| T51 | 348 | EOY23734.1 | *Theobroma cacao* | NAC domain class transcription factor, putative isoform 1 | 46% | 2.00E-16 |
| T52 | 348 | EOY23734.1 | *Theobroma cacao* | NAC domain class transcription factor, putative isoform 1 | 47% | 7.00E-18 |
| T53 | 297 | EOY23078.1 | *Theobroma cacao* | Phosphoenolpyruvate carboxylase-related kinase 1 isoform 4 | 71% | 8.00E-24 |

Table 1. Continued
| name | Length (bp) | Accession | Species | Putative function | Max Identity | E value |
| --- | --- | --- | --- | --- | --- | --- |
| T54 | 297 | EOY23078.1 | *Theobroma cacao* | Phosphoenolpyruvate carboxylase-related kinase 1 isoform 4 | 70% | 1.00E-22 |
| T55 | 348 | EOY23734.1 | *Theobroma cacao* | NAC domain class transcription factor, putative isoform 1 | 44% | 1.00E-17 |
| T57 | 326 | YP<br>004668894.1 | Myxococcus fulvus<br>HW-1 | amino acid<br>adenyltransferase>gb\|AEI67816.1\| | 43% | 5.4 |
| T58 | 331 | EOY29862.1 | *Theobroma cacao* | Uncharacterized protein TCM_037263 | 73% | 6.00E-25 |
| T59 | 331 | EOY29862.1 | *Theobroma cacao* | Uncharacterized protein TCM_037263 | 71% | 7.00E-21 |
| T60 | 330 | EOY29862.1 | *Theobroma cacao* | Uncharacterized protein TCM_037263 | 72% | 2.00E-21 |
| T61 | 328 | EOY29862.1 | *Theobroma cacao* | Uncharacterized protein TCM_037263 | 72% | 1.00E-22 |
| T62 | 331 | EOY29862.1 | *Theobroma cacao* | Uncharacterized protein TCM_037263 | 74% | 2.00E-25 |
| T63 | 90 | ADD09560.1 | *Trifolium repens* | unknown | 86% | 8.00E-08 |
| T64 | 47 | JX595015.1 | *Gossypium hirsutum* | Gossypium hirsutum clone NBRI_GE31150<br>microsatellite sequence | 92% | 1.00E-03 |
| T65 | 54 | AC190156.3 | *Pan troglodytes* | Pan troglodytes BAC clone CH251-487F13<br>from chromosome 7, complete sequence | 96% | 1.00E-15 |
| T66 | 59 | XM_00281433<br>9.1 | *Pongo abelii* | PREDICTED: Pongo abelii MCF.2 cell line<br>derived transforming sequence-like 2<br>(MCF2L2), partial mRNA | 84% | 0.6 |
| T67 | 256 | EOY26726.1 | *Theobroma cacao* | Alcohol dehydrogenase transcription factor<br>Myb/SANT-like family protein | 86% | 3.00E-24 |
| T68 | 143 | JX614796.1 | *Gossypium hirsutum* | Gossypium hirsutum clone NBRI_GE58446<br>microsatellite sequence | 82% | 5.00E-15 |
| T69 | 146 | JX614796.1 | *Gossypium hirsutum* | Gossypium hirsutum clone NBRI_GE58446<br>microsatellite sequence | 85% | 4.00E-17 |
| T72 | 19 | CP006252.1 | *Serratia liquefaciens* | Serratia liquefaciens ATCC 27592,<br>complete genome | 100% | 4.9 |
| T73 | 354 | EOX95883.1 | *Theobroma cacao* | Uncharacterized protein isoform 2 | 46% | 5.00E-13 |
| T74 | 170 | EOX95393.1 | *Theobroma cacao* | Class III peroxidase 70 | 84% | 1.00E-22 |
| T75 | 168 | AAQ56285.1 | *Oryza sativa*<br>*Japonica* | putative gag-pol protein | 80% | 2.00E-14 |

Table 1. Continued
| name | Length (bp) | Accession | Species | Putative function | Max Identity | E value |
| --- | --- | --- | --- | --- | --- | --- |
| T77 | 354 | EOY08498.1 | *Theobroma cacao* | RNA helicase family protein isoform 2 | 96% | 9.00E-67 |
| T78 | 354 | EOY08498.1 | *Theobroma cacao* | RNA helicase family protein isoform 2 | 95% | 6E-65 |
| T79 | 272 | WP_01023703<br>9.1 | *Citromicrobium bathyomarinum* | TonB-dependent receptor | 85% | 6.00E-05 |
| T80 | 254 | WP_00743912<br>3.1 | *Acetobacteraceae bacterium AT-5844* | Fis family transcriptional regulator | 60% | 5.3 |
| T81 | 272 | WP_01023703<br>9.1 | *Citromicrobium bathyomarinum* | TonB-dependent receptor | 85% | 6.00E-05 |
| T82 | 254 | AP010968.1 | *Kitasatospora* | Kitasatospora setae KM-6054 DNA, complete genome | 91% | 0.11 |
| T84 | 254 | AP010968.1 | *Kitasatospora* | Kitasatospora setae KM-6054 DNA, complete genome | 91% | 0.12 |
| T85 | 92 | EMJ14281.1 | *Prunus persica* | hypothetical protein PRUPE_ppa021229mg | 86% | 1.00E-07 |
| T86 | 91 | AC193213.3 | *Gallus gallus* | Gallus gallus BAC clone TAM31-41F20 from chromosome z, complete sequence | 90% | 0.0007 |
| T87 | 62 | JX615329.1 | *Gossypium hirsutum* | Gossypium hirsutum clone NBRI_GE59257 microsatellite equence | 84% | 7.00E-04 |
| T92 | 227 | EOY11709.1 | *Theobroma cacao* | Uncharacterized protein isoform 1 | 92% | 9.00E-30 |
| T93 | 70 | WP_01697191<br>7.1 | *Pseudomonas tolaasii* | hypothetical protein | 74% | 6.00E-04 |
| T94 | 84 | AAL06417.1 | *Glycine max* | reverse transcriptase | 76% | 4.00E-05 |
| T95 | 347 | EOY18193.1 | *Theobroma cacao* | UDP-XYL synthase 6 | 91% | 1.00E-35 |
| T96 | 347 | EOY18193.1 | *Theobroma cacao* | UDP-XYL synthase 6 | 91% | 3.00E-37 |
| T98 | 88 | AM447494.2 | *Vitis vinifera* | Vitis vinifera contig VV78X024983.11, whole genome shotgun sequence | 88% | 7.00E-10 |
| T99 | 45 | AC243111.1 | *Gossypium raimondii* | Gossypium raimondii clone GR_Ba0033O07-hvk, complete sequence | 90% | 3.9 |

### GO analysis of differentially expressed fragments

Blast2GO (http://www.blast2go.org/) was used to assigned GO IDs to the common genes based on the sequence of 86 differentially expressed fragments and performed Gene Ontology (GO) annotations to retrieve molecular function, biological process, and cellular component terms according to their putative function.

In terms of molecular function, these differentially expressed fragments were assigned to 7 functional groups, in which the binding was 8.24%, transferase activity was4.71%, catalytic activity was 2.35%, structural molecule activity, transcriptional regulation, antioxidant activity, and ubiquitin-protein ligase activity were 1.18% (Figure 1).

**Figure 1.**
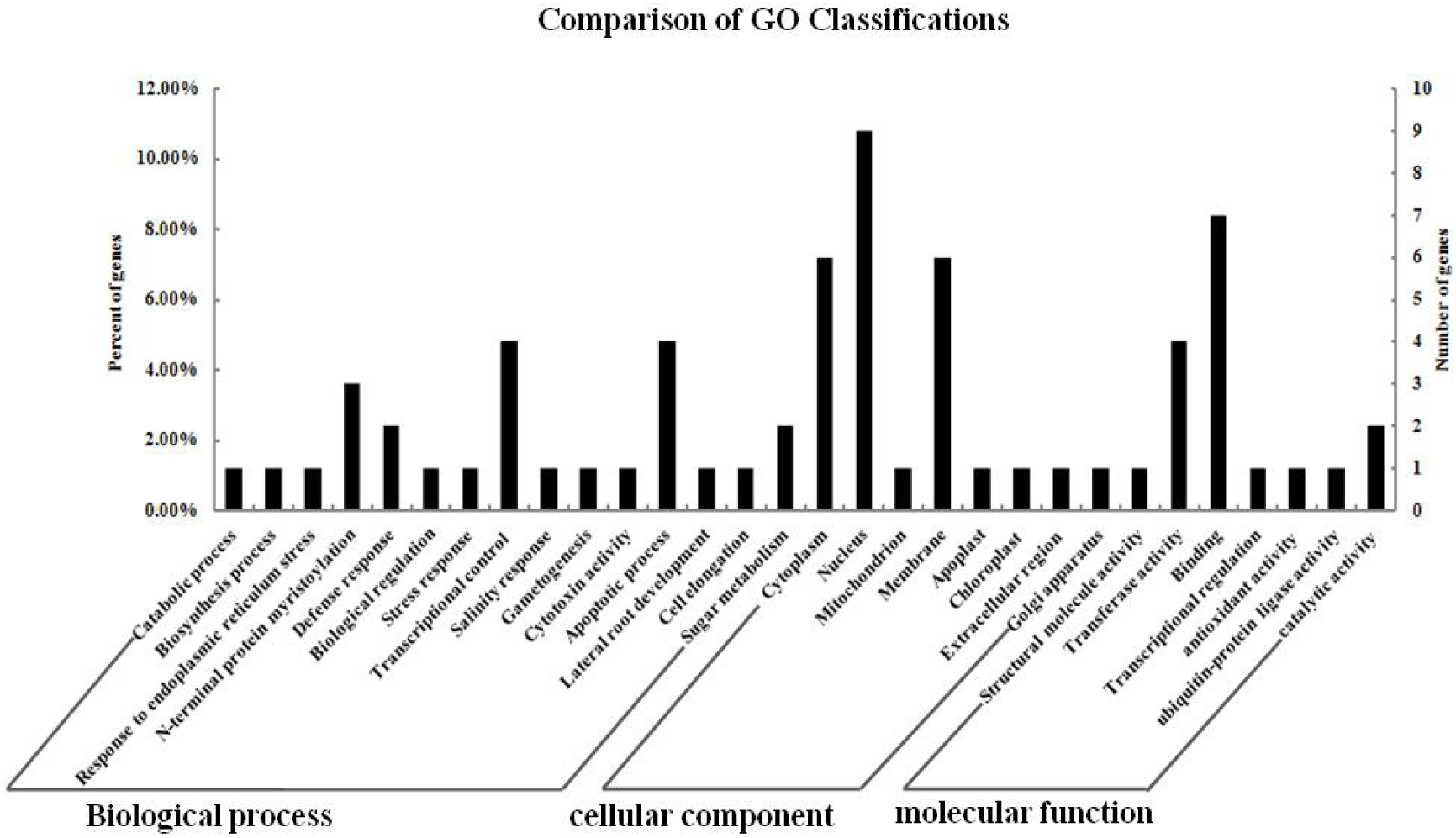
Gene Ontology classification of TDFs sequences on cDNA-AFLP.

In terms of biological process, these differentially expressed fragments were assigned to 15 functional groups, in which the transcriptional control and apoptotic process were 4.71%, N-terminal protein myristoylation was 3.53%, defense response and sugar metabolism were 2.35%, Catabolic process, biosynthesis process, response to endoplasmic reticulum stress, defense response, biological regulation, stress response, salinity response, gametogenesis, cytotoxin activity, lateral root development, cell elongation and were 1.18% (Figure 1).

In terms of cellular component, these differentially expressed fragments were assigned to 8 functional groups, in which the nucleus was 10.59%, cytoplasm and membrane were 7.06%, mitochondrion, apoplast, chloroplast, extracellular region, and Golgi apparatus were 1.18% (Figure 1).

### KEGG pathway mapping of differentially expressed fragments

Functional classification and pathway assignment of differentially expressed fragments were performed by KEGG. 86 differentially expressed fragments were assigned to 11 KEGG pathways (Table 2). The pathways with most representation by the differentially expressed fragments were beta-Alanine metabolism (3), RNA degradationrna (3), protein export (2), carbon metabolism (2), ribosome (1), regulation of actin cytoskeleton (1), oxidative phosphorylation (1), basal transcription factors (1), starch and sucrose metabolism (1), magnesium transporter (1), and plant-pathogen interaction (1).

**Table 2.** KEGG biochemical pathway analysis of TDFs sequences on cDNA - AFLP.

| Pathway name | Map No. | Count | Name |
| --- | --- | --- | --- |
| Protein export | 03060 | 2 | T1、 T90 |
| Ribosome | 03010 | 1 | T2 |
| Regulation of actin cytoskeleton | 04810 | 1 | T8 |
| beta-Alanine metabolism | 00410 | 3 | T12、 T26、 T27 |
| Oxidative phosphorylation | 00190 | 1 | T18 |
| RNA degradation | 03018 | 3 | T25、 T77、 T78 |
| Carbon metabolism | 01200 | 2 | T53、 T54 |
| Basal transcription factors | 03022 | 1 | T67 |
| Starch and sucrose metabolism | 00500 | 1 | T95 |
| magnesium transporter | 03284 | 1 | T28 |
| Plant-pathogen interaction | 04626 | 1 | T39 |

**Table 3.** Identification of differentially expressed proteins by MS/MS.

| SSP Numbers | Protein name | Accession no. | -10lgP | Molecular (kDa)/pI | Sequence coverage (%) | Peptides | *G.raimondii*.pepAccession no. |
| --- | --- | --- | --- | --- | --- | --- | --- |
| 0013 | ATP synthase D chain, mitochondrial [Theobroma cacao] | gb\|EOY18549.1 | 175.07 | 19.588/4.94 | 35 | 6 | Cotton_D_gene_10017606 |
| 1003 | Pathogenesis-related protein 10 (Gossypium hirsutum) | gb\|AAG18451.1\|AF305064_1 | 198.11 | 17.154/5.16 | 52 | 6 | Cotton_D_gene_10010317 |
| 1604 | ATP synthase CF1 beta subunit (chloroplast) [Gossypium raimondii] | ref\|YP_005087700.1\| | 189.58 | 46.501/5.96 | 20 | 6 | Cotton_D_gene_10029472 |
| 2005 | low molecular weight heat shock protein [Gossypium hirsutum] | gb\|ABW89469.1\| | 127.41 | 17.514/6.04 | 21 | 4 | Cotton_D_gene_10037266 |
| 2106 | glutathione S-transferase [Gossypium hirsutum] | gb\|ABW69659.2\| | 49.67 | 25.847/5.84 | 11 | 2 | Cotton_D_gene_10017216 |
| 2604 | enolase [Gossypium hirsutum] | gb\|ABW21688.1\| | 235.44 | 47.665/5.64 | 32 | 12 | Cotton_D_gene_10005023 |
| 3003 | Peroxisomal membrane 22 kDa (Mpv17/PMP22) family protein isoform 1 [Theobroma cacao] | gb\|EOY06118.1\| | 26.11 | 21.485/10.35 | 3 | 1 | Cotton_D_gene_10021212 |
| 3004 | MLP-like protein [Gossypium barbadense] | gb\|AER29900.1\| | 75.51 | 17.993/6.09 | 28 | 6 | Cotton_D_gene_10014373 |
| 4702 | ribulose-1,5-bisphosphate carboxylase/oxygenase large subunit (chloroplast) [Gossypium raimondii] | gb\|AFS41719.1\| | 178.73 | 53.894/6.57 | 25 | 12 | Cotton_D_gene_10018308 |
| 4713 | ATP synthase F1 subunit 1 (mitochondrion) [Gossypium hirsutum] | gb\|AFO69209.1\| | 120.18 | 51.587/8.05 | 6 | 3 | Cotton_D_gene_10036457 |
| 5720 | ribulose-1,5-bisphosphate carboxylase/oxygenase large subunit (chloroplast) [Gossypium raimondii] | gb\|AFS41719.1\| | 191.28 |  | 31 | 15 | Cotton_D_gene_10018308 |

## Proteomics analysis

### Protein expression profiles in Yamian A and Yamian B by 2-DE assay

Microspore abortion of YA-CMS occurred mainly between the stages of sporogenous cell and microsporocyte through the early-stage study of Cell morphological observation and comparison of physiology and biochemistry characteristics^(16)^. According to the bud developmental in cotton, the buds at the stages of sporogenous cell and microsporocyte in YA-CMS and YB were named A2, A3, B2, and B3 respectively. Thus, to further understand sterility mechanisms in YA-CMS, we performed a 2-DE analysis for the total protein of A2, A3, B2, and B3 (Figure S2). The total concentration of all detected protein spots was performed homogenization processing to obtain more accurate results. 1013, 1110, 1112 and 1110 protein spots were detected respectively in the 2-DE images of A2, B2, A3, and B3 by the PDQuest8.0.1 software. The molecular weights of these proteins ranged from 10 to 100 kDa, and the isoelectric point from 3.0 to 10.0.

A total of 11 protein spots changed significantly (P < 0.05) in relative abundance by a minimum of a 2.0-fold change in at least one stage between YA-CMS and YB through the point-to-point comparison and statistical analysis. Most of these differential spots displayed quantitative changes, but some displayed qualitative changes. Eight protein spots of them had significant quantitative differences in expression between YA-CMS and YB. For examples, 2604 spot was up-regulated, whereas 3004 was down-regulated in flower buds from the sporogenous cell stage of the YA-CMS plants, but instead in the YB plants. 0013, 2005, and 3003 spots were up-regulated whereas 1003, 2106, and 4713 were down-regulated in flower buds from the microsporocyte stage of the YA-CMS plants but instead in the YB plants (Figure 2). There were three protein spots that had significant qualitative differences in expression between YA-CMS and YB. For example, 1604 and 4702 were expressed only in flower buds from sporogenous cell and microsporocyte stage of the YB plants but not in YA-CMS plants. 5720 was detected only in flower buds from the sporogenous cell and the microsporocyte stage of the YA-CMS plants but not in YB plants (Figure 2).

**Figure 2.**
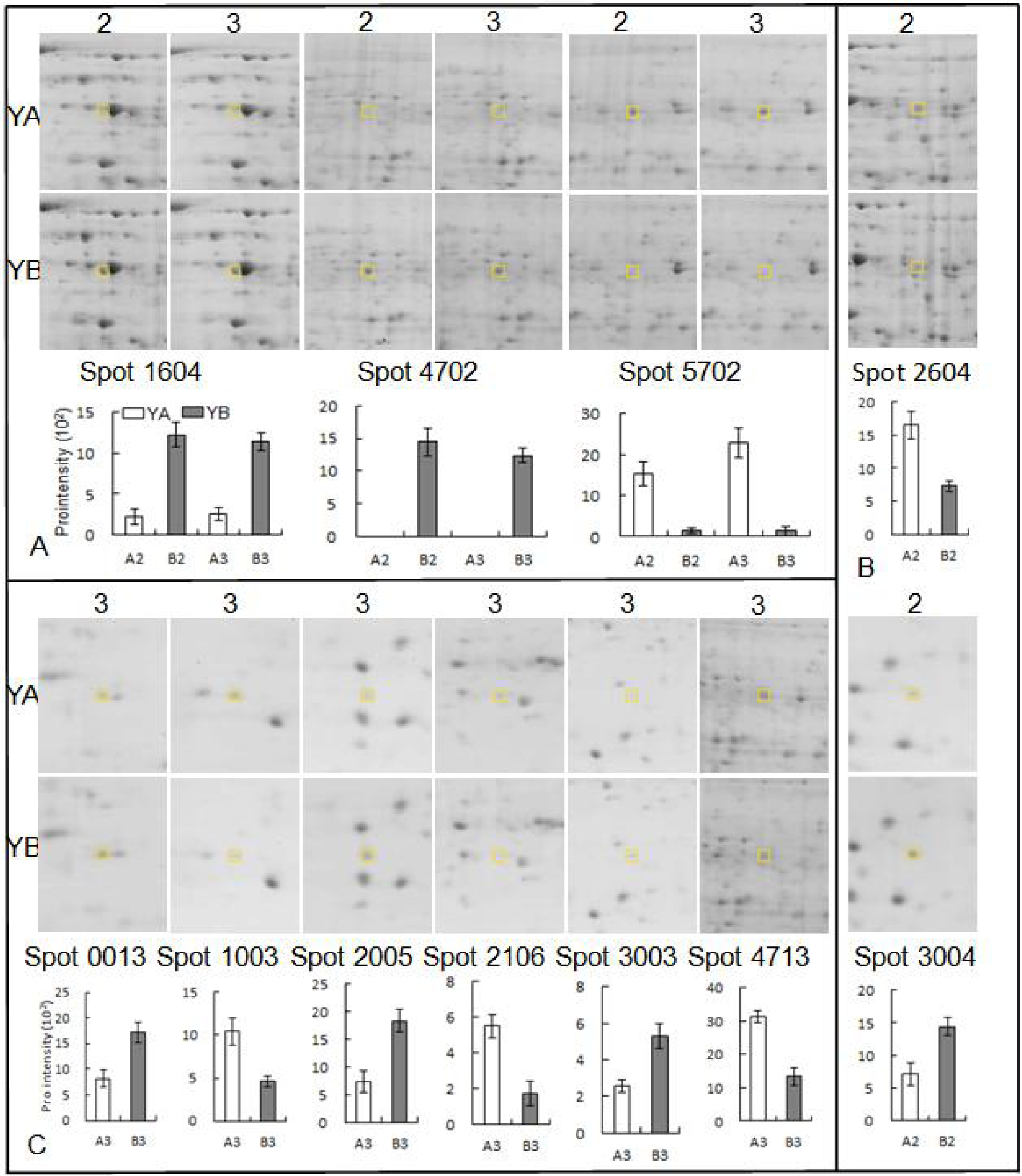
Enlarged differentially expressed protein spots from flower buds between YA-CMS and YB. A, B, C: represent differentially expressed proteins in sporogonium and microsporocyte stage, sporogonium stage, microsporocyte stag from flower buds.

### Identification and functional annotation of differentially expressed proteins

All 11 differentially expressed spots were analyzed by LC-Chip-ESI-QTOF-MS. 11 spots were successfully identified by MASCOT and PEAKS 6.0 Software’s searched protein database NCBInr green plants and cotton. The 11 spots included ATP synthase D chain (spot 0013), Pathogenesis-related protein 10 (spot 1003), ATP synthase CF1 beta subunit (spot 1604), low green plants and cotton. The 11 spots included ATP synthase D chain (spot 0013), Pathogenesis-related protein 10 (spot 1003), ATP synthase CF1 beta subunit (spot 1604), low molecular weight heat shock protein (spot 2005), glutathione S-transferase (spot 2106), enolase (spotmolecular weight heat shock protein (spot 2005), glutathione S-transferase (spot 2106), enolase (spot 2604), Peroxisomal membrane 22 kDa (Mpv17/PMP22) family protein isoform 1 (spot 3003), MLP-like protein (spot 3004), ribulose-1,5-bisphosphate carboxylase/oxygenase large subunit (spot 4702), ATP synthase F1 subunit 1 (spot 4713) and ribulose-1,5-bisphosphate carboxylase/oxygenase large subunit (spot 5720) (**Tab1e 3**).

Gene Ontology (GO) annotations to retrieve molecular function, biological process, and cellular component terms according to their putative function. Blast2GO (http://www.blast2go.org/) was used to assign GO IDs to the 11 differentially expressed proteins (Figure 3). Spot 0013 participated in the creature process of photorespiration, had zinc ion binding function, and was located in the mitochondrion. Spot 1003 participated in the creature process of mRNA modification and were located in the membrane. Spot 2005 participated in the creature process of protein folding and was located in the cytoplasm. Spot 2106 participated in the proteasomal ubiquitin-dependent protein catabolic process and had glutathione transferase activity. Spot 2604 responded to light stimulus and had DNA binding. Spot 4713 participated in ATP biosynthetic process which was located in the mitochondrion. Other differentially expressed proteins were respectively in mitochondrion, plastid, and nucleus (Figure 3).

**Figure 3.**
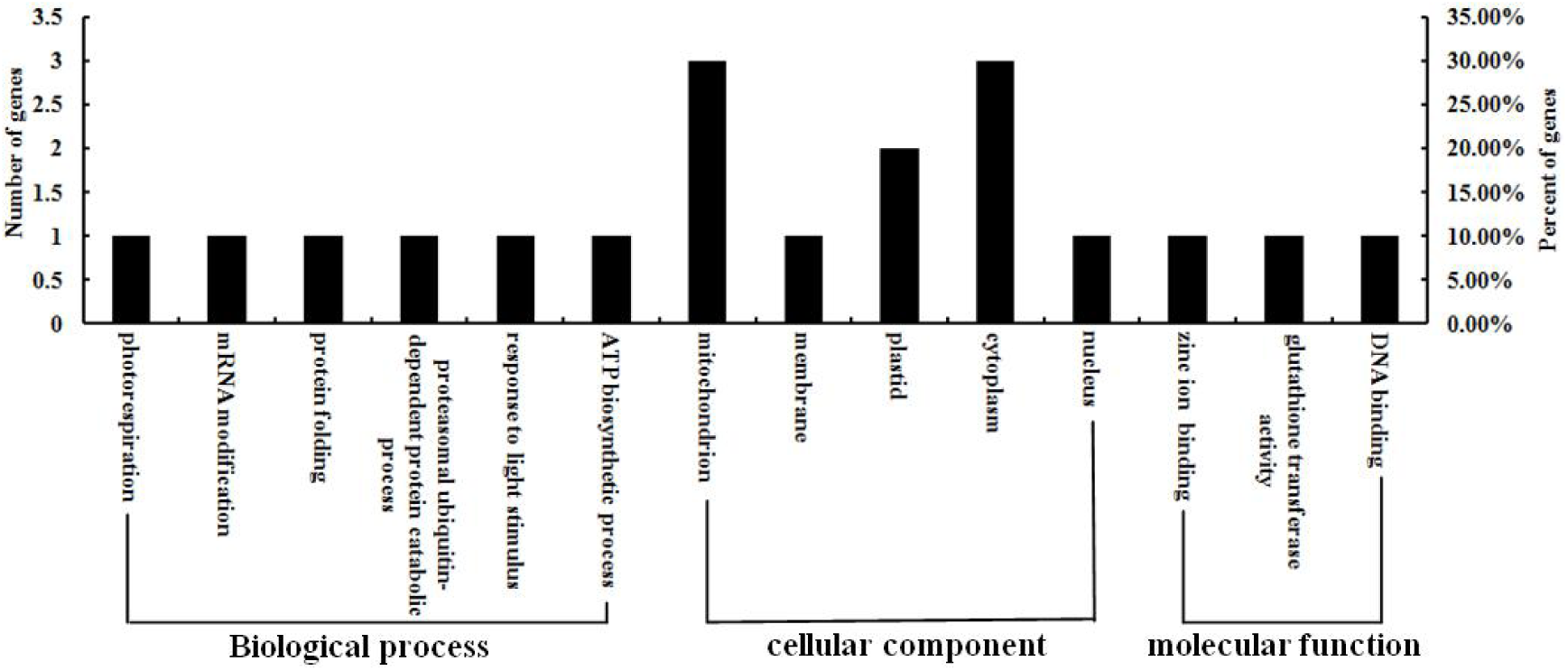
Gene Ontology classification of differentially expressed proteins.

The differential protein information about protein functions in the metabolic pathway was revealed by KEGG pathway enrichment analysis. KEGG analysis showed that six differentially expressed proteins were significantly enriched in the five identified classes. These proteins were involved in multiple biological processes, including carbon fixation in photosynthetic organisms, glyoxylate and dicarboxylate metabolism, glycolysis/gluconeogenesis, photosynthesis, and oxidative phosphorylation (Table 4).

**Table 4.**
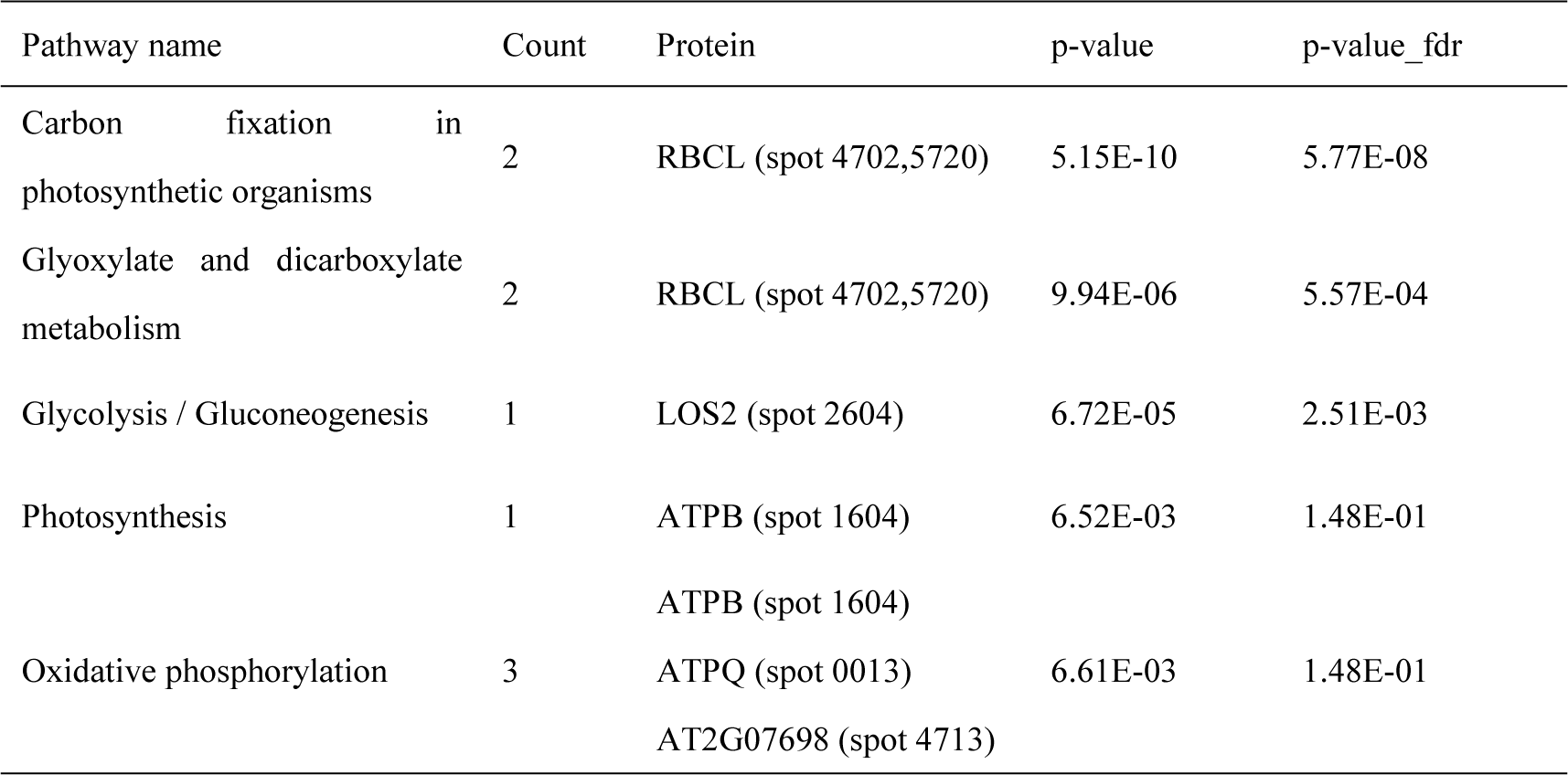
Enriched KEGG pathway-based sets of differentially expressed proteins.

The insights into the functional mechanisms of living cells can be provided by protein-protein interaction networks. 11 proteins were recognized as key nodes with various relationships in biological interaction networks by using the online tools of STRING 9.05 (Figure 4). Five proteins were mainly involved in the carbohydrate metabolism (ATPQ[spot 0013], ATPB[spot 1604], LOS2[spot 2604], RBCL[spot 4702/5720] and AT2G07698[spot 4713]).

**Figure 4.**
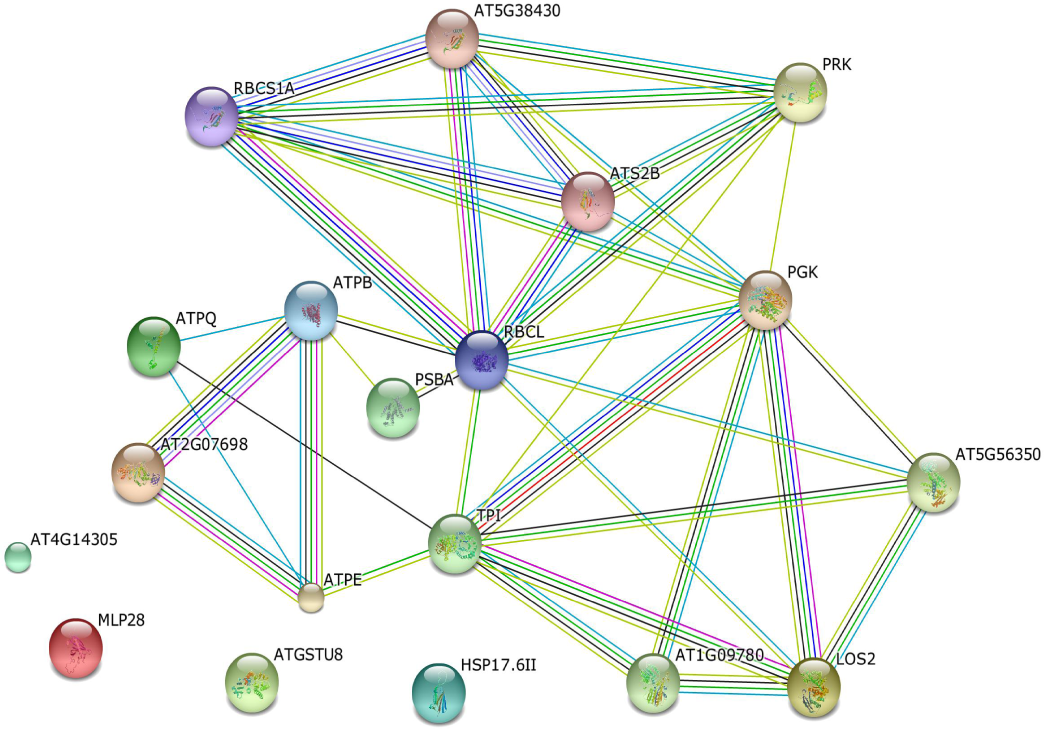
Biological interaction network of the differentially expressed proteins. Blue indicates co-occurrence evidence; red indicates fusion evidence, green indicates neighborhood evidence, yellow indicates text-mining evidence, purple indicates experimental evidence, light blue indicates database evidence and black indicates coexpression evidence.

### Validation of genes and proteins of differential abundance

To verify the differential abundance of expression gene come from cDNA-AFLP, seven genes were selected to perform qRT-PCR using the equal amounts of cDNA templates from the buds of seven different development periods respectively of both the fertile and sterile plants. The results observed by qRT-PCR were the same as that of cDNA-AFLP (Figure S3 **and** Figure 5). T75 and T74 were both detected at the 2nd stage of flower buds on Yamian A. T12 was detected at the 2nd, 3rd and 4th stage of flower buds on Yamian A, which was the peak period of microspore abortion, and not detected in the flower buds of other periods on Yamian A and all periods on Yamian B. T26 and T39 were respectively detected at the 6th, 7th and 5th, 6th, 7th stage of flower buds on Yamian A, which was after microspore abortion, and not detected in the flower buds of other periods on Yamian A and all periods on Yamian B. T67 and T81 was detected at the 2nd, 3rd and 4th stage of flower buds on Yamian B, and not detected in the flower buds of other periods on Yamian B and all periods on Yamian A. These results suggested that the genes from cDNA-AFLP could stable expression.

**Figure 5.**
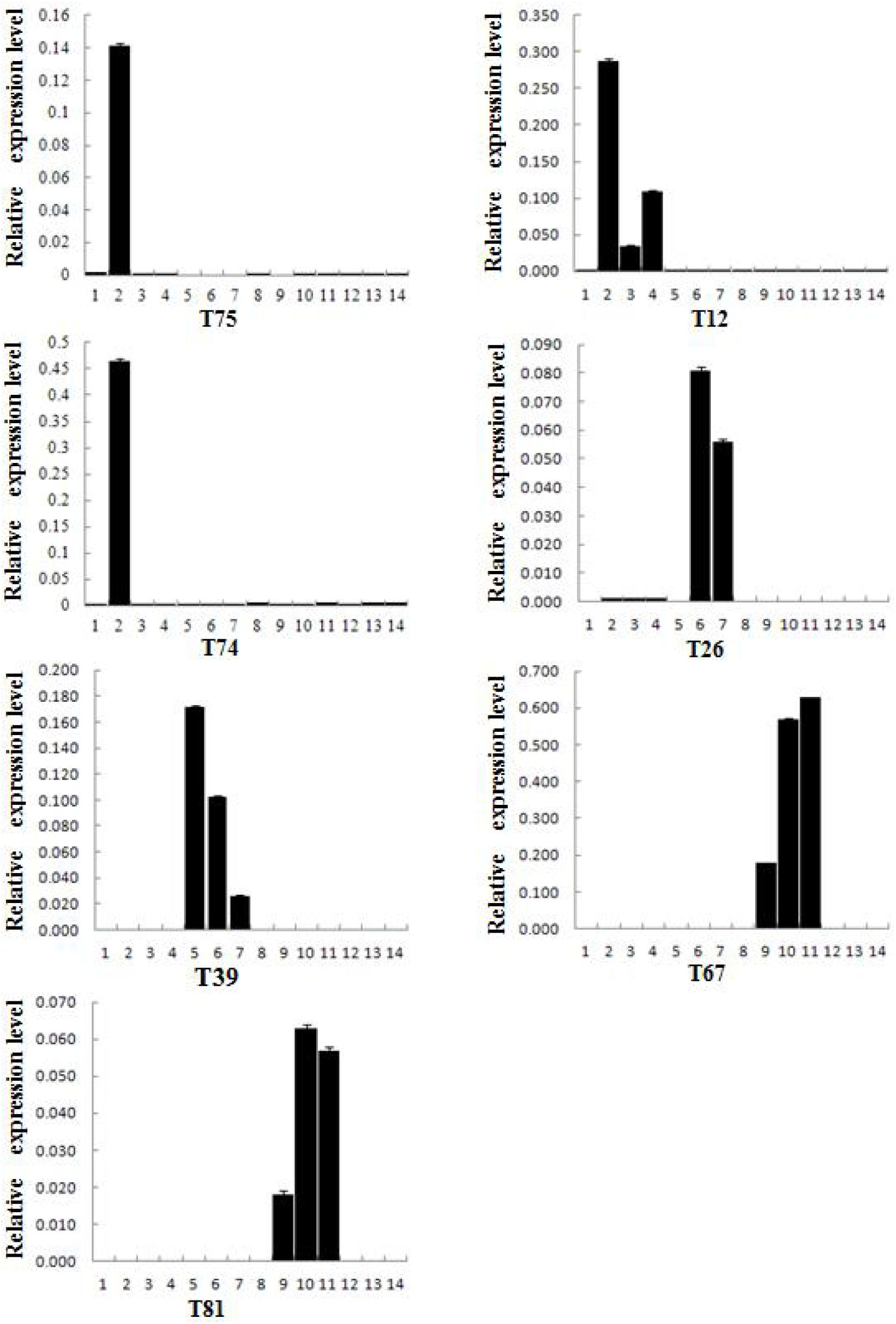
Validation of TDFs sequences on cDNA - AFLP by qRT-PCR. 1-7 and 8-14 respectively represent seven different development periods of the bud on Yamian A and Yamian B.

Seven coding genes that corresponded to differentially expressed proteins were selected to analyze in the mRNA expression levels by qRT-PCR to examine our 2-DE results and verify the differences in protein abundance at the transcriptional level (Figure 6). The expression level of 1003 at the 2nd, and the 7th stages of the floral buds in Yamian A was lower compared to that at the same stages in the Yamian B, but instead at the 3^rd^ stage of the flower bud, and was not great difference at other periods between Yamian A and Yamian B. The expression level of 3004 at the 2nd stage of the flower bud in Yamian A was lower compared to that at the same stages in the Yamian B, but instead at the 3rd stage of the flower bud. The expression level of 0013, 1604, 4702 and 2005 at the 2nd, 3^rd^ stage of the flower buds in Yamian A was lower compared to that at the same stages in the Yamian B. The expression level of 4713 at the 2nd, 3rd stage of the flower buds in Yamian A was higher compared to that at the same stages in the Yamian B. To study the abortive cause of YA-CMS, we used a transcriptomics and proteomics approach to compare the differentially expressed transcripts and proteins respectively in the buds of before, middle and after microspore abortion stage and the key stage of pollen abortion (Stage2, 3) between YA-CMS and its maintainer YB. In the analysis of the transcriptome, we acquired 99 TDFs; at the same time, we obtained 11 differential expression protein spots during the study of proteomics. We found that the results of functional annotation, GO and KEGG analyses of some TDFs were the same or similar with the corresponding analysis results of some differential expression protein spots, while others were quite different. These results indicated that two perspectives study on the transcriptomics and proteomics came with a high degree of consistency and complementarity. We made a comprehensive study and more just decision which may be the main cause on the anther abortive of YA–CMS by conjoint analysis of the transcriptomics and proteomics, which is the major finding of this study.

**Figure 6.**
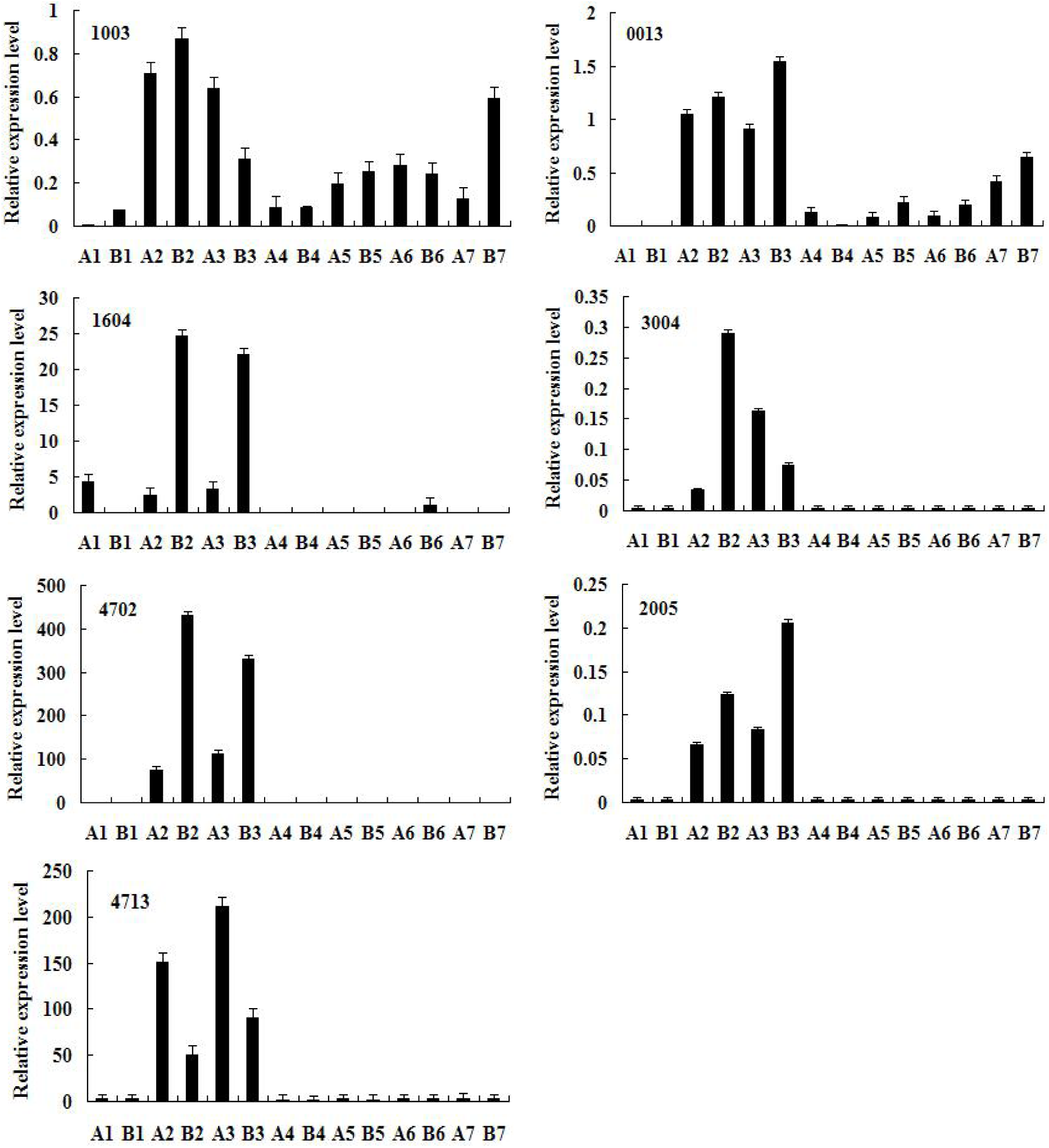
Real-PCR validation of different abundances of proteins at the mRNA level. 1-7 and 8-14 respectively represent seven different development periods of the bud on Yamian A and Yamian B.

### The consistency of transcriptomics and proteomics

ATP synthase is the key enzyme in the process of mitochondria oxidative phosphorylation. Mitochondria ATP synthase belongs to the energy storage “F” type, which consists of two parts, Fo and F1 regions. Fo region is located within the inner membrane of plant mitochondria and functions as a proton channel. F1 is the active enzyme center and composed of alpha, beta, and so on subunits. The binding sites of beta subunits have the activity of catalytic ATP synthesis or hydrolysis^(20)^.

Pollen development is a process of high energy consumption, if some or some gene products interfere with the function of mitochondrial FoF1-ATP synthase, which may lead to the abortion of pollen^(21)^. Many studies have shown that ATP synthase has a close relation with cytoplasmic male sterility. For example, Li et al. ’s study of protein interactions on chili pepper profiled that the decrease of the activity and amount of ATP synthase affected the development of pollen, and thus caused the cytoplasmic male sterility^(22)^. The use of the SNP marker of ATP synthase gene could simply, rapidly and easily identify the cytoplasmic male sterile line CMS-D8^(23)^. The study results of *atp1*^(24)^, *atp4*^(25)^, *atp6*^(26)^, *atp8*^(27)^ and *atp9*^(28)^ by researchers showed that these genes may be related to cytoplasmic male sterility in plants. Studies have also found that 29 mtDNA regions associated with CMS have been identified, and these recombinant chimeric genes were involved in the promoter region and part of the coding region of the ATP synthase subunit gene. Kong et al. ’s RNA editing analysis of ATP synthase genes on the cotton CMS line H276A and maintainer line and restorers line showed forty-one RNA editing sites, and two new stop codons were detected and suggested the ATP synthase genes might be an indirect cause of cotton CMS^(29)^. Proteins analysis in CMS of Wheat showed ATP synthases could be associated with abnormal pollen grain formation and male sterility^(30)^.

In this study, We found 1 ATPase-like protein (TDF 15) from the transcriptome, and 3 ATP synthase subunits from the proteome, 1 ATP synthase beta subunit (1604-2, 1604-3), 1 ATP synthase D chain (0013-3) and 1 ATP synthase F1 subunit 1 (4713-3). Among them, the ATP synthase beta subunit was significantly up-regulated in the sporogenous cell and microsporocyte stage of the YB plants but not in YA-CMS plants at the same stages. The expression of the ATP synthase D chain was significantly reduced in the microsporocyte stage of the YA-CMS plants compared with the YB plants in the same period. ATP synthase F1 subunit 1 was more significantly up-regulated in the microsporocyte stage of the YA-CMS plants than in YB plants. ATP synthase-like protein gene was significantly up-regulated in the sporogenous cell and microsporocyte stage of the YA-CMS plants but not in YB plants at the same stages. Zheng et al. ’s study found that ATP synthase beta subunit and ATP synthase D chain were down-regulated in Male Sterile Mutant YX-1 anthers of Wolfberry^(31)^. Li et al. ’s study found that ATP synthase beta subunit was not expressed in the wheat BNS male sterile line, but expressed in its transformation line^(32)^. These results were consistent with ours. The differential proteomics was studied with the upland cotton cytoplasmic sterile line 104-7A, maintainer line, and restorer line by Xu Qi, and the result found that ATP synthase beta subunit was expressed only in the restorer line, while without expression in sterile and maintainer lines^(33)^. Wu et al. ’s study found that ATP synthase D chain was down-regulated in *Capsicum annuum* L. CMS anthers, but ATP synthase beta subunit was up-regulated in the same material^(34)^. These results were not consistent with ours. According to previous ultrastructure observations, the sporogenous cell and microsporocyte of the YA-CMS plants both contained numerous abnormal mitochondria. The above results showed that the down-regulated of ATP synthase beta subunit and ATP synthase D chain led to internal energy metabolism disorder, caused large mitochondria abnormal disintegration, then affected the development of anther, ultimately the male sterility occurred in the YA-CMS plant. Also, the disagreements with the up and down-regulated of ATP synthase and its subunit from different male sterile lines in the different plants, even the same kind of plant but different genotypes may be caused by their self-different abortion mechanisms, these different sterility mechanisms are still not well understood and needs further research.

## The complementarity of transcriptomics and proteomics

### The relationship between the pollen abortion and the differences of UDP-glucosyltransferase (UGT) in the YA-CMS and in YB plants

UDP-glycosyltransferases are the major glycosyl transferase in plants. They can transfer the glycosyl groups of the activated donor molecule (mainly uridine diphosphate glucose) to the receptor molecule (such as secondary metabolites such as flavonoids, phytohormones such as cytokinins, herbicides, insecticides, etc.), thus regulating the location of the receptor molecule in the cell and its biological activities such as solubility and transport in organisms ^(35, 36)^. UGT plays an important role in regulating glycosylation and energy storage of secondary metabolites in organisms, endogenous hormone activity and toxicity relief of exogenous toxins^(37, 38)^. In this study, two different primers (E2M7 and E15M3) were used to amplify the difference of the highest consistency with uridine diphosphate glucosyltransferase gene in the buds of the YA-CMS plant during the peak period of abortion (sporogenous cell proliferation stage, microspore mother cell stage and meiosis stage of microspore development). The allogenic fragments were not amplified in the maintainers of the same period. This suggests that the uridine diphosphate glucosyltransferase gene may play a role in the peak period of microspore abortion of cotton male sterile line Yamian A, and may be related to the microspore abortion of Yamian A, but this hypothesis still needs further experiments to verify.

### The relationship between the pollen abortion and the differences of ribosomal protein in the YA-CMS and in YB plants

The ribosome is a protein-nucleic acid complex enzyme system^(39)^. As the main site of protein synthesis in cells, the integrity of ribosome structure and the coordination of the quantity of each component are the necessary conditions to ensure the effective and correct synthesis of protein^(40)^. Although it is generally believed that these ribosomal proteins play an important role in protein synthesis, more and more ribosomal proteins have been reported to have many other functions. For example, they can play a role in the regulation of cell apoptosis, proliferation, development, and malignant transformation by participating in transcription, RNA processing, DNA repair and replication^(38)^. Zhou et al. found that the ribosome proteins were essential for anther development and male sterility in sterile buds when they studied genetic male sterile line ‘AB01’ in Chinese cabbage^(5)^. This study indicates that there is a certain relationship between ribosomal protein and plant male sterility.

A gene fragment with 95% consistency with the 60S ribosomal protein L13a of castor was isolated, and expressed only at the peak of microspore abortion of cotton cytoplasmic male sterile line Yamian A (i.e., sporogenous cell proliferation stage, microspore mother cell stage and meiosis stage of microspore development), but no expression was observed at the microspore development other periods of male sterile line Yamian A and the whole anther formation period of the maintainer line. This result suggests that the 60S ribosomal protein L13a gene may be involved in the development of Microspore in male sterile line, which is related to the abortion of Microspore in male sterile lines, but this hypothesis still needs further experiments to verify.

### The relationship between the pollen abortion and the differences of NAC transcription factor in the YA-CMS and YB plants

NAC transcription factors are one of the largest families of transcription factors peculiar to plants. They have many functions. They are widely involved in the formation of lateral roots, secondary walls, shoot apical meristem, senescence and flowering of plants, as well as the response to abiotic and biological stresses^(41)^. Chen et al. found that the 9 NAC transcription factor genes were downregulation expressed and 6 NAC transcription factor genes were up-regulation expressed in sterile buds when they studied CMS in Wucai^(42)^.

Our study results showed that the gene fragment T51, which was 46% consistent with cocoa NAC transcription factors, was amplified from the mixed buds of microspore development tetrad, monocyte and binucleate pollen grains and mature pollen grains of cotton maintainer line by using selection primers E7M2, but not amplified in the buds of sterile line in corresponding period. In terms of cell morphology, At this stage, the pollen sac of male sterile line anthers contracted and became smaller after microspore mother cells disintegrated completely. Then the tapetum cells elongated radially and filled the pollen sac during the tetrad formation of fertile anthers, and finally formed pollen sac without pollen grains. This result suggests that pollen abortion of CMS lines may be caused by mutation or silence of the NAC transcription factor gene, but this hypothesis still needs further experimental verification.

### The relationship between the pollen abortion and the differences of ribulose-1,5-bisphosphate carboxylase/oxygenase in the YA-CMS and YB plants

Ribulose-1,5-bisphosphate carboxylase/oxygenase (RUBISCO) is widely distributed in the organelles of photosynthesis. It is a key enzyme for fixing CO2 in plant photosynthesis and also participates in the photorespiration pathway of plants. Riboketose-1,5-diphosphate carboxylase/oxygenase is composed of 8 small subunits (12-18 kD) encoded by nuclear genes and 8 large subunits (50-60 kD) encoded by chloroplasts. The small subunits have regulatory functions, and the enzyme activity locates on the large subunits. Kurepa J and Smalle J A found that the oxidative stress caused by promoting the generation of superoxide anion induced the formation of covalently linked ribulose-1,5-bisphosphate carboxylase/oxygenase large subunit dimer, and its formation coincided with the loss of chloroplast function when they studied the effects of oxidative stress on tobacco^(43)^.

Current studies on Cytoplasmic Male Sterility in many plants have shown that RUBISCO is related to fertility. Chen et al. showed that the expression of the RUBISCO subunit in two stages of wheat cytoplasmic-nuclear interaction male sterile line was significantly down-regulated, and this result suggested that energy metabolism might be closely related to anther development^(44)^. Liu et al. found that the activity of RuBp carboxylase in cytoplasmic male sterile lines of maize, sorghum, rice, wheat, and tobacco was higher than that in their corresponding maintainer lines, indicating that there was a certain relationship between RuBp carboxylase and cytoplasmic male sterility in plants^(45)^. Ren Yan also identified five RUBISCO or its large subunits in the differential proteome analysis of anthers of double Recessive Genic Male Sterile Lines and fertile lines of *Gossypium hirsutum* Linn^(46)^.

Two ribulose-1,5-bisphosphate carboxylase/oxygenase subunits were found to be associated with CMS in chloroplasts. The spots on the 2-DE diagram show that the molecular weight is the same, but the isoelectric point is not the same, one is acidic, the other is alkaline. The acidic large subunit was expressed only in the critical period of abortion of maintainer line, while the alkaline large subunit was only expressed in the sterile line at the critical period of abortion. The difference of two ribulose-1,5-bisphosphate carboxylase/oxygenase subunits in chloroplast between sterile lines and maintainer lines may be caused by the reactive oxygen species in different degrees, and this may be related to anther fertility of cytoplasmic male sterility, and the specific mechanism needs further study.

### The relationship between the pollen abortion and the differences of low molecular weight heat shock protein in the YA-CMS and YB plants

Heat shock proteins (HSPs) are a kind of stress proteins induced and synthesized by organisms under the influence of adverse environmental factors such as high temperature, hypoxia, starvation, and heavy metal ions. They can improve the heat resistance of cells and have the functions of molecular chaperone and regulation. At present, heat shock proteins have become the focus of molecular biology research, and there are some reports on male sterility. Heat shock protein *HSP70* gene transcription was blocked in sterile line, which made cell meiosis not normal, resulted in the number of anther mitochondria in sterile line, and then the pollen development could not get enough energy, resulted in pollen abortion^(47, 48)^. Zeng et al. also found heat shock protein 22kDa in anther differential proteomics of soybean cytoplasmic male sterile line NJCM2A, and speculated that it might lead to abnormal mitochondrial development, thus resulting in inadequate energy supply for pollen development and eventually abortion^(49)^. Su et al. found that six BoHSP70 genes were highly expressed in the binuclear-pollen-stage buds of a male fertile line compared with its near-isogenic sterile line when they studied the HSP70 family genes in cabbage^(50)^.

In this study, low molecular weight heat shock protein (SSP2005) was found in the buds of infertile line A and maintainer line B during the critical period of abortion, and its expression in maintainer line was higher than that in male-sterile line B. our results were similar to the result of Su et al. The difference of expression of low molecular weight heat shock protein (SSP2005) between sterile lines and maintainer lines indicates that SSP2005 may be related to anther fertility of cytoplasmic male sterility.

Combined with all results of the transcriptome, proteome and early cytological, physiological and biochemical studies of cytoplasmic male sterile line Yamian A and its maintainer line Yamian B in cotton, We speculated and constructed a schematic diagram, which implied that there might be a connection among UGT, *NAC TFs*, ATP, RUBISCO, heat shock protein, Peroxidase to cytoplasmic male-sterile of Yamian A (Figure 7). However, the occurrence of cytoplasmic male sterility has certain temporal and spatial specificity. Further studies are still needed to determine the exact nature of the full mechanism of cytoplasmic male sterility in Yamian A.

**Figure 7.**
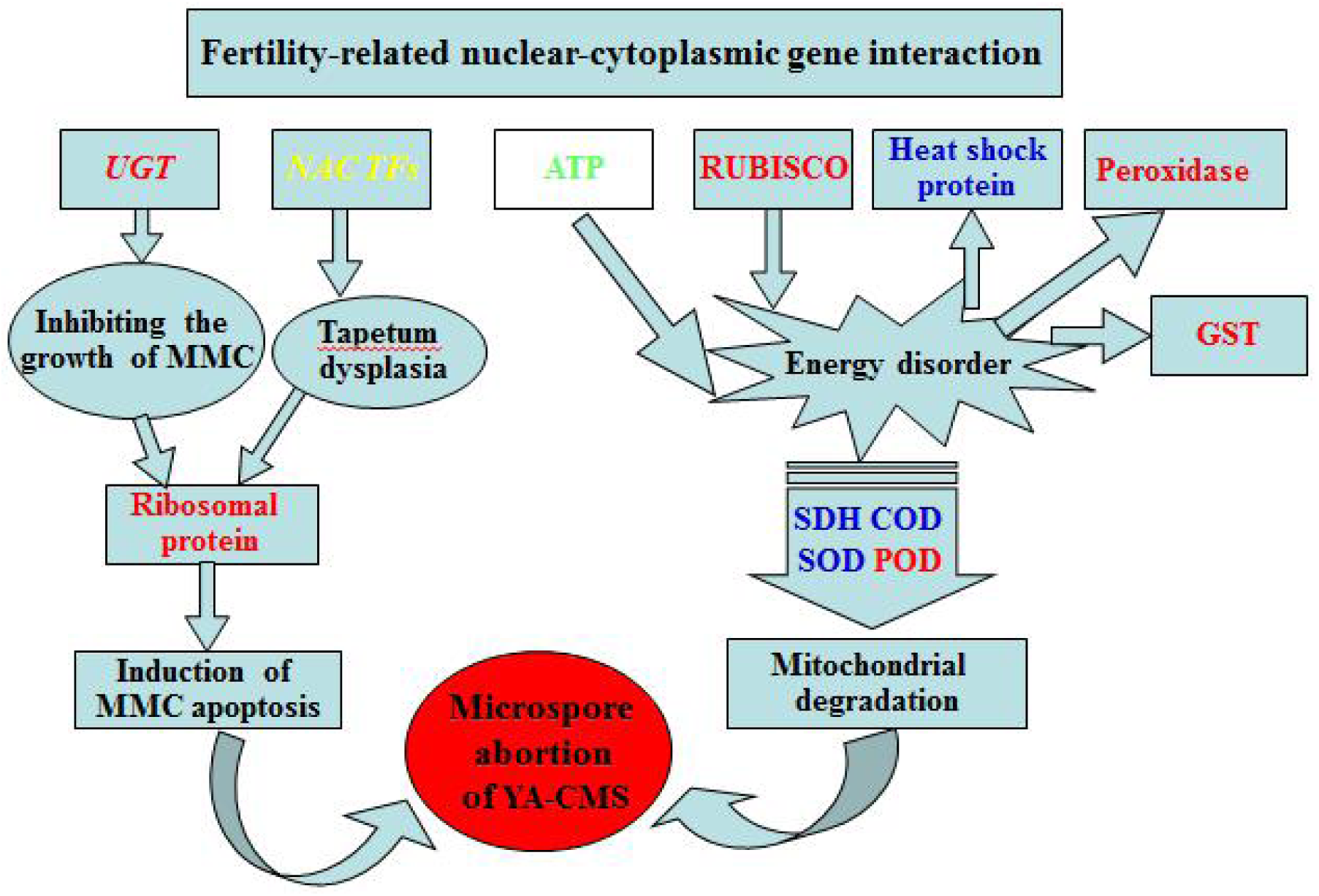
Hypothetical interaction network of microspore abortion in cytoplasmic male sterile line Yamian A. Red font: up-regulated expression; Blue font: downward expression; Yellow font: no expression; Green Font: Up and Down Expressions.

## ABBREVIATIONS

CMS: cytoplasmic male sterility
DEGs: differentially expressed genes
DEPs: differentially-expressed proteins
UGT: UDP-glucosyltransferase
RUBISCO: ribulose-1,5-bisphosphate carboxylase/oxygenase
GST: glutathione S-transferase
IMS: induced male sterility
GMS: genic male sterility
ROS: reactive oxygen species
YA-CMS: CMS line Yamian A
YB: Yamian B
qRT-PCR: quantitative real-time PCR
cDNA-AFLP: cDNA amplified fragment length polymorphism
GO: gene ontology
KEGG: kyoto encyclopedia of genes and genomes
TDFs: transcript-derived fragments
HSPs: heat shock proteins

## AUTHOR INFORMATION

### Notes

The authors declare no competing financial interest.

## Author Contributions

JH and HZ conceived and designed the experiments. HZ and JW performed the experiments and analyzed the data. JH provided the Yamian A and Yamian B materials. HZ and JW analyze the data. HZ wrote the manuscript. YQ, RP, RM and FL revised the manuscript. JH critically revised the manuscript and supervised the writing.

## ACKNOWLEDGEMENT

This project was supported by National Key R&D Program of China Grant 2018YFD0100301, Shanxi Key R&D Program Grant 201703D221002-4, 201703D221007-11 to JH, and Anyang Scientific and Technological Project Grant 96 to HZ.

## SUPPLEMENTARY MATERIALS

The supporting information includes Figure S1-S3 and Table S1-S3, which corresponds to the supplementary documents.

**Figure S1.**
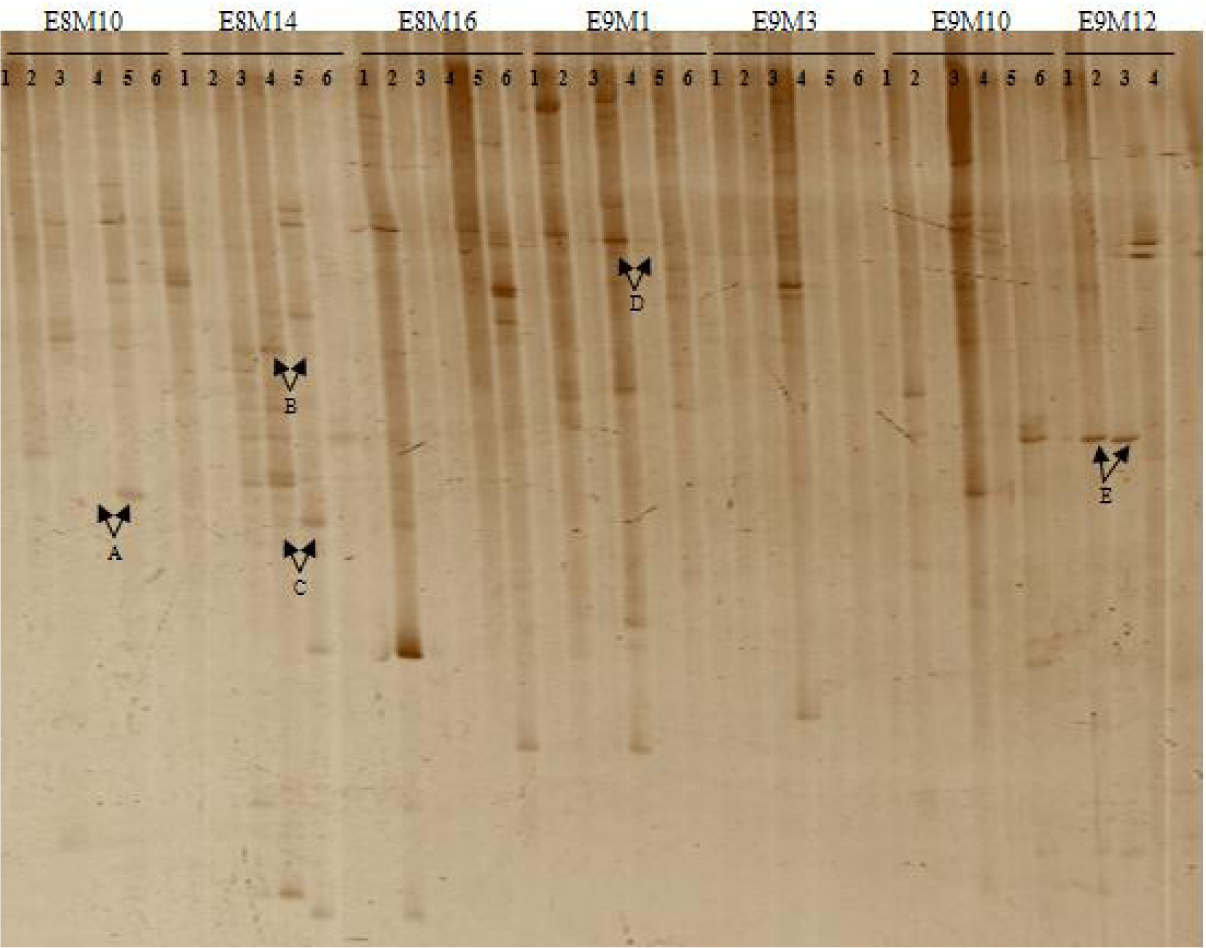
Selective amplified products of several primer pairs. 1, 3, 5 and 2, 4, 6 represent anther of before, middle and after microspore abortion stage of Yamian A and Yamian B. A, B, C, D, E represents different kinds of expressed bands between Yamian A and Yamian B.

**Figure S2.**
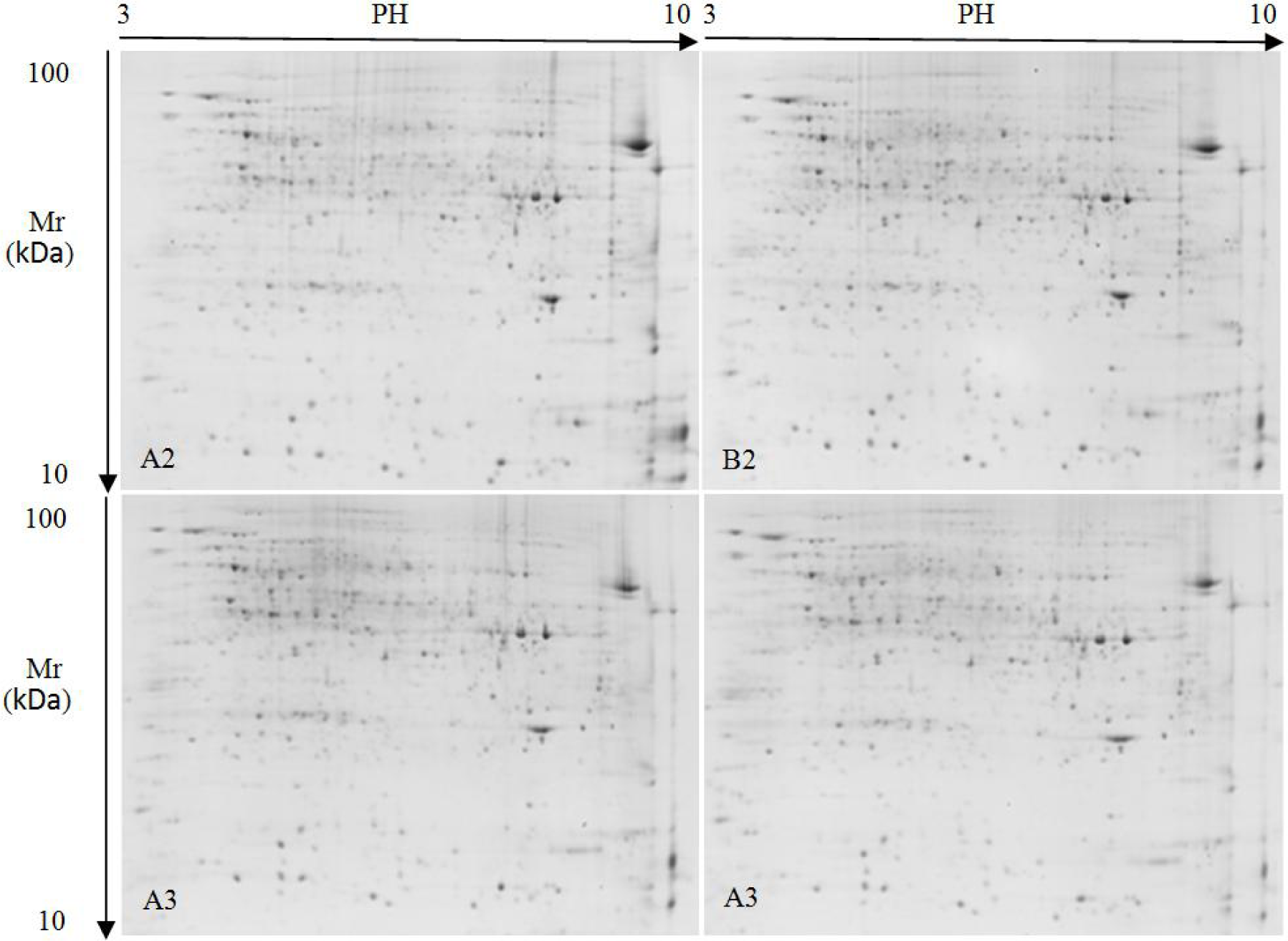
2-DE images of flower buds proteins in sporogenous cell and microsporocyte stage from the YA-CMS and YB. A2: sporogenous cell stage of YA-CMS; A3: microsporocyte stage of YA-CMS; B2: sporogenous cell stage of YB; B3: microsporocyte stage of YB.

**Figure S3.**
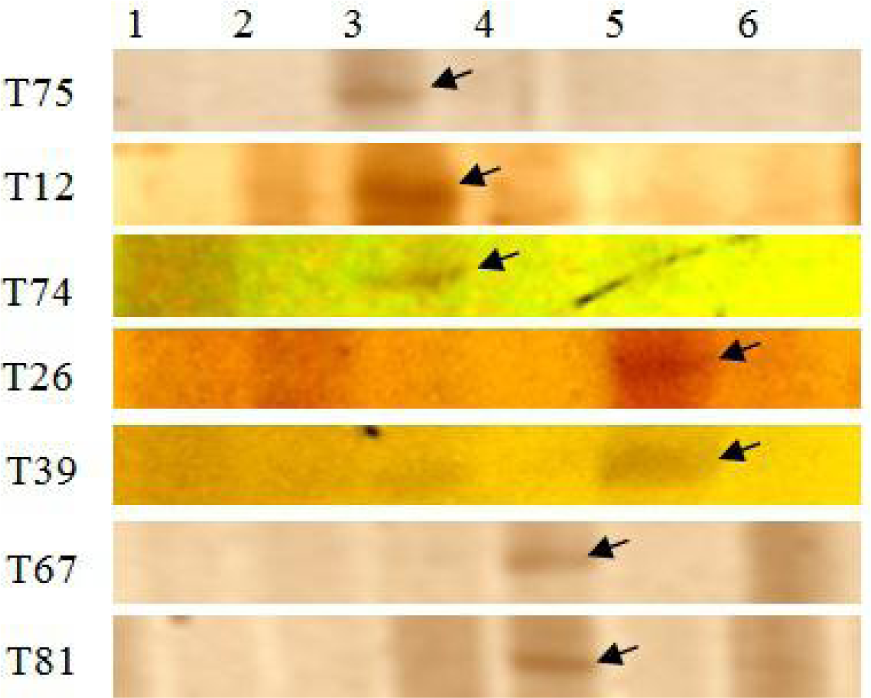
Parts of TDFs on cDNA - AFLP. 1, 3, 5, and 2, 4, 6 represent anther of before, middle and after microspore abortion stage of Yamian A and Yamian B.

**Table S1.** Bud developmental stages in cotton^(16)^.

| No. | Stage | Bud transverse diameter (BTD) (mm) |
| --- | --- | --- |
| 1 | Sporogonium stage | BTD≤1.50 |
| 2 | Sporogenous cells stage | 1.50<BTD≤2.16 |
| 3 | Microsporocyte stage | 2.16<BTD≤2.60 |
| 4 | Meiosis stage | 2.60<BTD≤4.60 |
| 5 | Tetrad stage | 4.60<BTD≤5.90 |
| 6 | mononuclear and binucleate pollens stage | 5.90<BTD≤9.93 |
| 7 | Pollen maturation stage | BTD>9.93 |

**Table S2.**
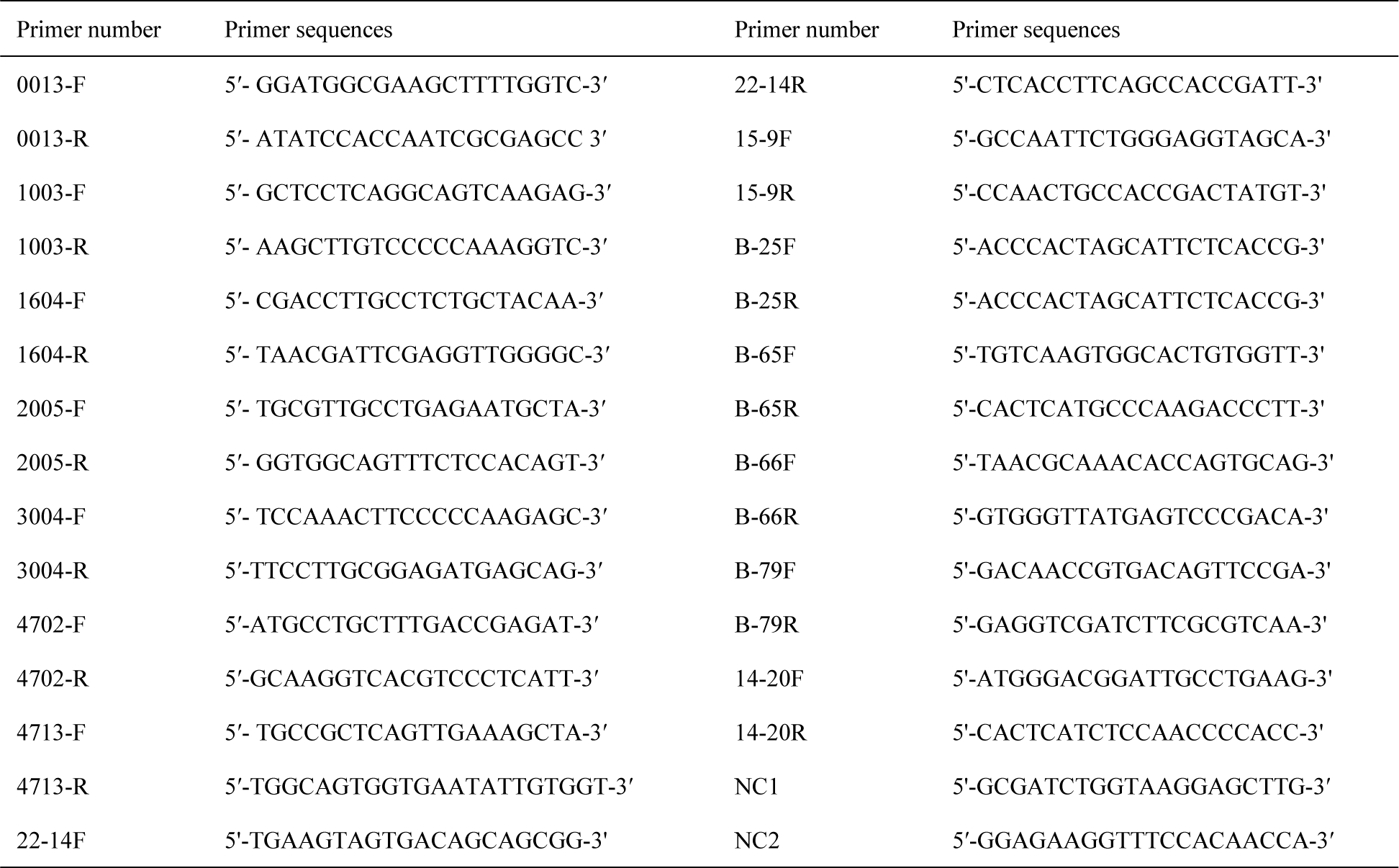
Primers used in qRT-PCR.

**Table S3.** Important type of gene differential expression.

| Type | S1 | F1 | S2 | F2 | S3 | F3 | Total bands | primer number |
| --- | --- | --- | --- | --- | --- | --- | --- | --- |
| 1 | + | - | - | - | - | - | 12 | 8 |
| 2 | - | - | + | - | - | - | 100 | 36 |
| 3 | - | - | - | - | + | - | 42 | 17 |
| 4 | - | + | - | - | - | - | 17 | 8 |
| 5 | - | - | - | + | - | - | 20 | 5 |
| 6 | - | - | - | - | - | + | 5 | 4 |
| 7 | + | - | + | - | - | - | 12 | 4 |
| 8 | - | - | + | - | + | - | 63 | 11 |
| 9 | + | - | - | - | + | - | 20 | 6 |
| 10 | - | + | - | - | - | + | 7 | 1 |
| 11 | - | + | - | + | - | - | 14 | 3 |
| 12 | - | - | + | + | - | - | 20 | 6 |
| 13 | - | - | + | + | + | - | 17 | 3 |
| 14 | + | + | + | - | - | + | 5 | 1 |
| 15 | + | + | + | - | - | + | 15 | 1 |

S1, S2, S3 and F1, F2, F3 represent anther of before, middle and after microspore abortion stage of Yamian A and Yamian B respectively; +: with a band, -: no band.

